# Cofilin promotes actin turnover and flexibility to drive coordinated cell movements *in vivo*

**DOI:** 10.1101/2024.12.17.628979

**Authors:** Ana Maria do Carmo, Ji Hong Sayo, Rodrigo Fernandez-Gonzalez

## Abstract

Embryos display a striking ability to repair wounds rapidly, with no inflammation or scarring. Embryonic wound healing is driven by the collective movement of the cells adjacent to the wound. The cells at the wound edge polarize actin and the molecular motor non-muscle myosin II, forming a supracellular cable around the wound that generates force and coordinates cell movements to close the lesion. Actin network contraction has been associated with the disassembly of the actin filaments that form the network. We found that the actin-severing protein Cofilin and its co-factor Aip1 accumulated at the edge of epidermal wounds in *Drosophila* embryos. Reducing Cofilin activity or levels slowed down wound closure, indicating that Cofilin is necessary for rapid wound healing. Using quantitative microscopy, we showed that Cofilin controls F-actin turnover at the wound edge, but not F-actin polarity or contractile force generation. Combining genetic and pharmacological manipulations, we found that F-actin turnover at the wound edge must be tightly regulated for wounds to close rapidly. Computational modelling suggested that Cofilin may contribute to rapid wound repair by maintaining a flexible actin network around the wound. Consistent with this model, fluorescence fluctuation analysis revealed that F-actin networks at the wound edge were significantly more rigid when we reduced Cofilin activity. Together, our results indicate that Cofilin promotes F-actin turnover at the wound margin to maintain a flexible actin network and facilitate rapid contraction and wound healing.

## Introduction

Wound healing in embryos occurs rapidly, without inflammation or scarring (Longaker et al., 1990; Rowlatt, 1979). Embryonic wound repair is a conserved morphogenetic process driven by the collective migration of the cells around the wound. Upon wounding, actin and the molecular motor non-muscle myosin II become polarized in the cells immediately adjacent to the wound, forming a supracellular cable at the wound edge (Kiehart et al., 2000; Martin & Lewis, 1992; McCluskey et al., 1993; Wood et al., 2002). The actomyosin cable contracts, coordinating cell movements to draw the wound closed (Abreu-Blanco et al., 2012; Fernandez-Gonzalez & Zallen, 2013; Kobb et al., 2017; Wood et al., 2002; Zulueta-Coarasa & Fernandez-Gonzalez, 2018).

Actin filament disassembly is critical for actin network contraction. Stabilization of actin filaments during cytokinesis in budding yeast slows down the scission of the two daughter cells (Mendes Pinto et al., 2012). Similarly, F-actin severing is key for the contraction of the actin networks that drive cellularization of the *Drosophila* embryo (Xue & Sokac, 2016), gastrulation (Jodoin et al., 2015), or wound healing in pupal tissues (Antunes et al., 2013). Consistent with a role for actin disassembly in embryonic wound closure, F-actin levels at the wound edge decrease as the wound repairs (Fernandez-Gonzalez & Zallen, 2013), and F-actin stabilization slows down wound healing (Kobb et al., 2019).

Cofilin is one of the main actin-severing proteins (Bamburg & Bernstein, 2010). Cofilin binds preferentially to the pointed end of actin filaments (Carlier et al., 1997), where monomer removal dominates. Cofilin binding induces a change in filament conformation that increases the flexibility of the Cofilin-decorated segment (Brannon et al., 2011; McCullough et al., 2008). Thus, Cofilin binding creates a boundary between the flexible (Cofilin-decorated) and the rigid (non-decorated) ends of the filament that renders the filament susceptible to severing when thermal fluctuations occur (Elam et al., 2013; McCullough et al., 2008; Prochniewicz et al., 2005). The actin-binding protein AIP1 is recruited by Cofilin and is directly involved in the severing process, serving as a clamp that disrupts the contacts between actin filament subunits (Oosterheert et al., 2025). Cofilin is regulated by phosphorylation: the serine/threonine kinase LIM domain kinase (LIMK) phosphorylates Cofilin and inhibits actin binding, thus inactivating Cofilin (Agnew et al., 1995; Arber et al., 1998). Whether Cofilin regulates actin dynamics during embryonic wound healing is currently unknown.

## Results

### Cofilin is necessary for rapid embryonic wound healing

To investigate if Cofilin may be implicated in embryonic wound healing, we examined the localization of an endogenously-tagged form of Cofilin (Buszczak et al., 2007; Ikawa & Sugimura, 2018) during wound repair in embryos of the fruit fly *Drosophila melanogaster*. In the intact epidermis, Cofilin localized predominantly to the nuclei of epidermal cells, with an additional cytosolic pool (Fig. 1A). Strikingly, there was little-to-no accumulation of Cofilin at cell-cell junctions. Upon wounding, Cofilin accumulated at the edge of the wound, with Cofilin levels at the wound margin increasing by 11 ± 1% (mean ± standard deviation, SD) (Fig. 1A-B). A GFP-tagged form of the Cofilin cofactor AIP1 (Buszczak et al., 2007) was also largely absent from cell junctions in the intact epidermis, but similar to Cofilin, AIP1 accumulated at the wound edge by 27 ± 12%. (Fig. 1C-D). Thus, our results suggest that Cofilin may control actin dynamics to drive rapid embryonic wound repair.

**Figure 1.**
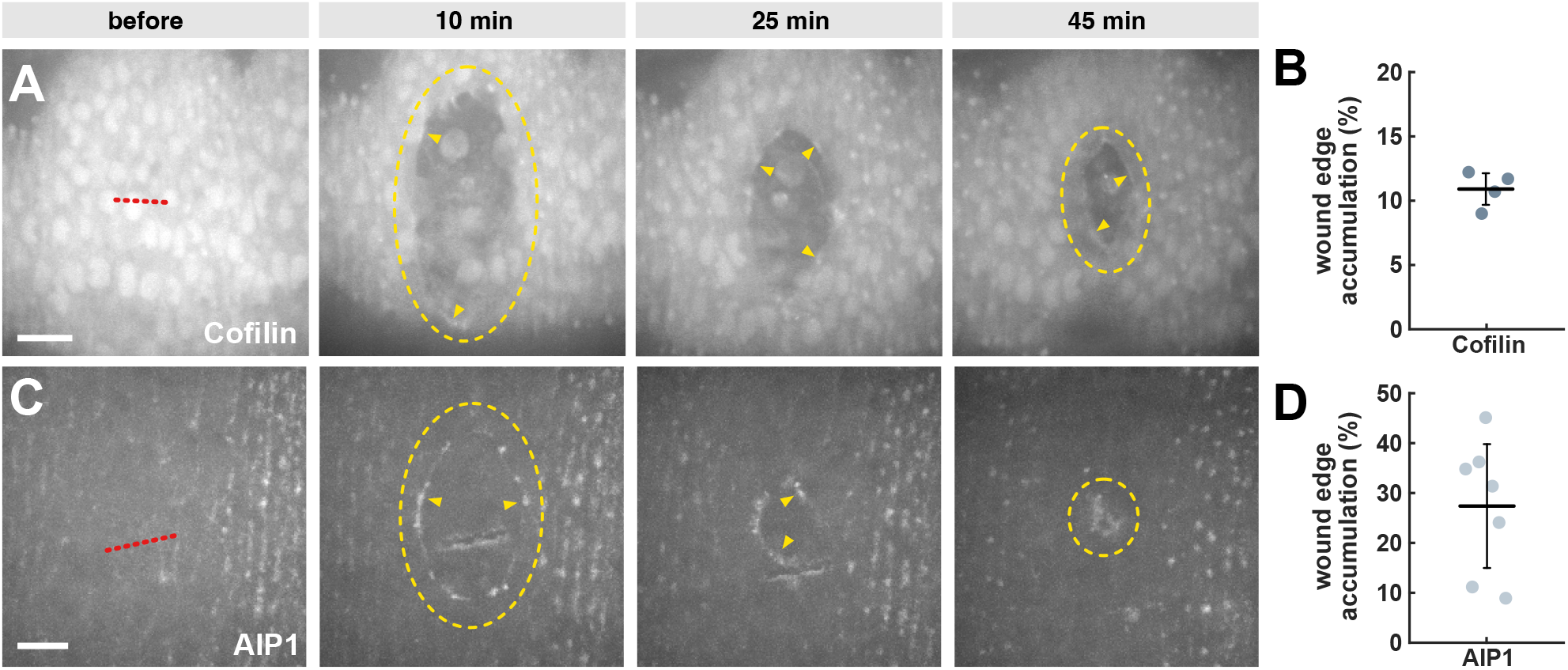
Cofilin and its cofactor AIP1 both localize to the wound edge. **(A, C)** Wound closure in embryos expressing endo-Cofilin:GFP (A) or endo-AIP1:GFP (C). Arrowheads indicate fluorescence accumulation at the wound edge. Red dotted lines indicate wound sites, yellow dashes outline the wounds. Time is with respect to wounding. Anterior left, ventral down. Bars, 10 μm. **(B, D)** Maximum fluorescence accumulation at the wound edge for endo-Cofilin:GFP (B, *n* = 4 wounds in as many embryos) or endo-AIP1:GFP (D, *n* = 7 wounds) Error bars, SD; line, mean.

Cofilin is necessary for cellularization of the *Drosophila* embryo (Xue & Sokac, 2016) and for gastrulation (Jodoin et al., 2015). Thus, to investigate if Cofilin is necessary for rapid embryonic wound healing, we established a method to reduce Cofilin activity while bypassing early embryonic development. We used the UAS-Gal4 system (Brand & Perrimon, 1993) to overexpress LIMK after gastrulation, using *daughterless-Gal4* (Wodarz et al., 1995) as the driver. To test if LIMK overexpression reduced Cofilin activity, we fixed control embryos and embryos overexpressing LIMK 10-11 hours after egg laying, and we stained the embryos with antibodies against total Cofilin and phospho-Cofilin (P-Cofilin), the inactive form of Cofilin (Xue & Sokac, 2016) (Fig. S1A-B). LIMK overexpression resulted in a twofold increase in the ratio of P-Cofilin to total Cofilin (*P* = 0.04, Fig. S1C). Thus, our results indicate that LIMK overexpression inactivates Cofilin in the *Drosophila* embryonic epidermis.

To determine if Cofilin contributes to rapid embryonic wound healing, we quantified the dynamics of wound closure in embryos overexpressing LIMK (Fig. 2A-B, Video S1). Embryos also expressed GFP:UtrophinABD (Rauzi et al., 2010), a fluorescent F-actin reporter. Reduced Cofilin activity was associated with a slower wound healing response, as demonstrated by a slower reduction of the wound perimeter over time (Fig 2C). To quantify the delay in wound repair, we fitted exponential decays to the wound perimeter *vs*. time curves and used the decay constant as a metric for the rate of wound healing (Hunter et al., 2018). We found that reducing Cofilin activity slowed down wound closure by 26% (*P* = 0.02, Fig. 2D). We obtained similar results when we quantified wound healing dynamics in heterozygous *cofilin* mutants (*tsr* ^*+/-*^), in which the rate of wound healing decreased by 39% (*P* = 0.005, Figure S2A-D). Together, our results show that Cofilin activity is necessary for rapid embryonic wound repair.

**Figure 2.**
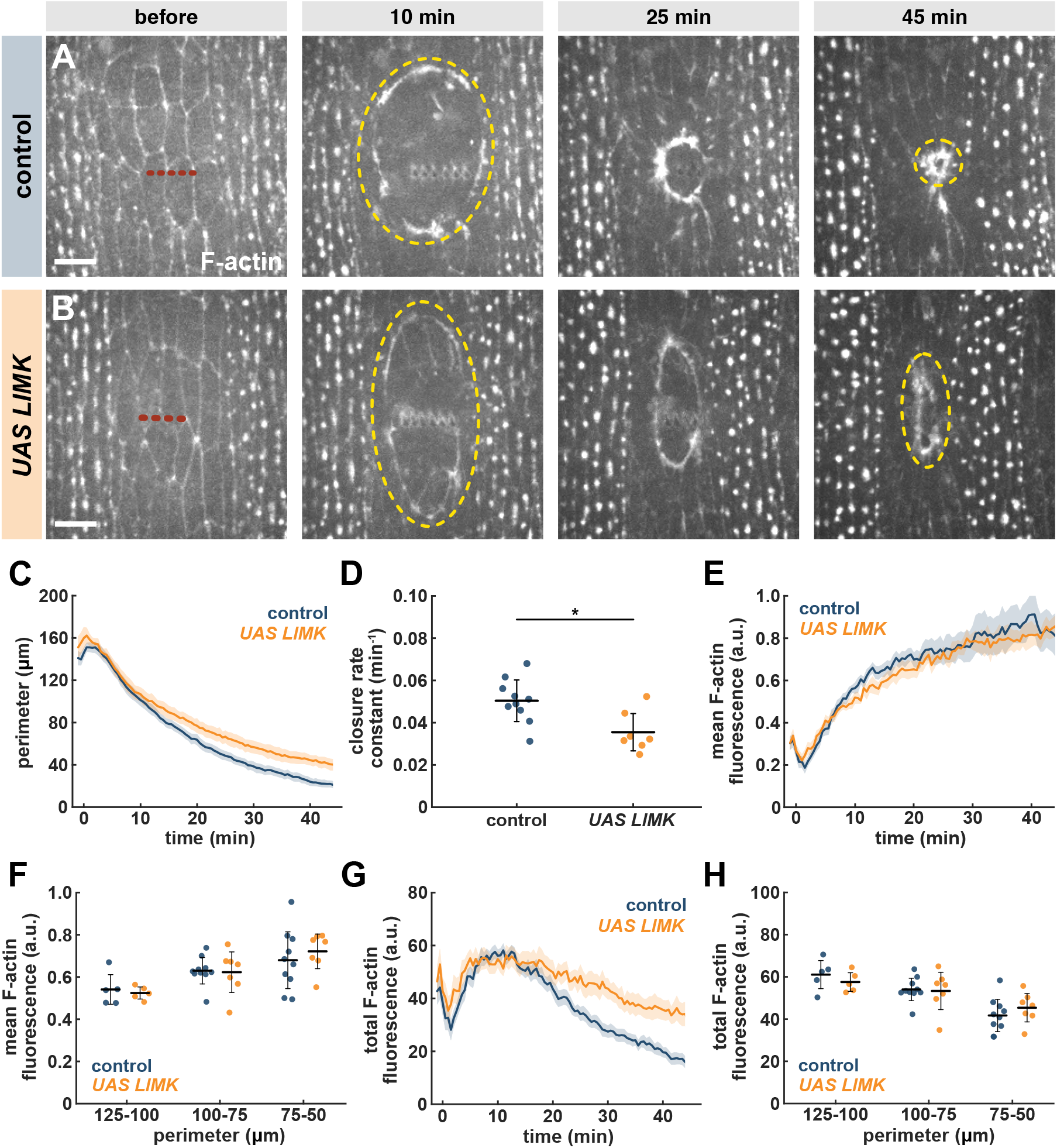
Cofilin activity is necessary for rapid wound closure. **(A-B)** Wound closure in control (A) and LIMK-overexpressing (B) embryos expressing GFP:UtrophinABD. Red dotted lines indicate wound sites, yellow dashes outline the wounds. Time is with respect to wounding. Anterior left, ventral down. Bars, 10 μm. **(C-H)** Wound perimeter over time (C), wound closure rate constant (D), mean or total F-actin fluorescence at the wound edge over time (E or G, respectively), or embryo averages binned by perimeter (F or H, respectively) in control (blue, *n* = 10 wounds) or LIMK-overexpressing embryos (orange, *n* = 7). (C, E, G) Error bars, standard error of the mean (SEM). (D, F, H) Error bars, SD; line, mean. * *P* < 0.05 (Mann-Whitney test).

### Cofilin promotes F-actin turnover at the wound edge

To investigate the mechanisms by which Cofilin contributes to rapid wound healing, we quantified F-actin fluorescence around the wound. Mean F-actin levels at the wound edge increased 3.4 ± 2.4-fold in controls over the first 45 min after wounding (*P* = 0.0002, Fig. 2A, E). Reducing Cofilin activity did not affect the polarization of F-actin to the wound edge, with F-actin levels increasing 3.1 ± 1.4-fold in LIMK-overexpressing embryos (*P* = 0.0006, Fig. 2B, E). The polarization of actin to the wound edge remained unaffected by reducing Cofilin activity when we compared wounds of similar perimeters (Fig. 2F). We obtained similar results when we measured F-actin polarization in *tsr* ^*+/-*^ mutants, in which F-actin accumulation around the wound was not disrupted (Fig. S2A-B, E-F). Similarly, mean myosin levels at the wound edge in embryos expressing a GFP-tagged form of the myosin regulatory light chain (encoded for by the *sqh* gene in *Drosophila*) (Royou et al., 2004) were not affected by reducing Cofilin activity (Fig. S3A-D). In contrast, total F-actin at the wound edge decreased faster in control embryos than in embryos with reduced Cofilin activity (Fig. 2G) or levels (Fig. S2G). However, the difference in total F-actin levels disappeared when we compared wounds of similar perimeter (Figs. 2H and S2H). Overall, our results suggest that Cofilin is dispensable for actomyosin polarization to the wound edge.

Cofilin flexibilizes and severs actin filaments, and thus, Cofilin could regulate the dynamics of F-actin at the wound edge. To investigate this possibility, we used fluorescence recovery after photobleaching (FRAP) of the F-actin reporter GFP:UtrophinABD (Fig. 3A). We previously showed that the FRAP dynamics of GFP:UtrophinABD match the dynamics of other F-actin and monomeric actin reporters (Kobb et al., 2019). We photobleached a 2.7 x 2.7 μm^2^ region on the actin cable around wounds that had closed to 50% of their maximum area. There were no significant differences in the mean fluorescence signal prior to photobleaching (Fig. 3B). We found that the degree of fluorescence recovery was lower in embryos with reduced Cofilin activity when compared to controls (Fig. 3C): the percent of initial signal recovered (the mobile fraction) decreased by 25% in LIMK-overexpressing embryos (47 ± 15% *vs*. 62 ± 19%, respectively, *P* = 0.04, Fig. 3D). Similarly, in *tsr* ^*+/-*^ mutants, the F-actin mobile fraction decreased by 43% with respect to controls (34 ± 3% *vs*. 59± 17%, respectively, *P* = 0.009, Fig. S4A-D). Reducing Cofilin activity or levels did not affect the rate of fluorescence recovery (the half-time, Figs. 3E and S4E). Thus, our experiments suggest that Cofilin promotes F-actin turnover during embryonic wound healing.

**Figure 3.**
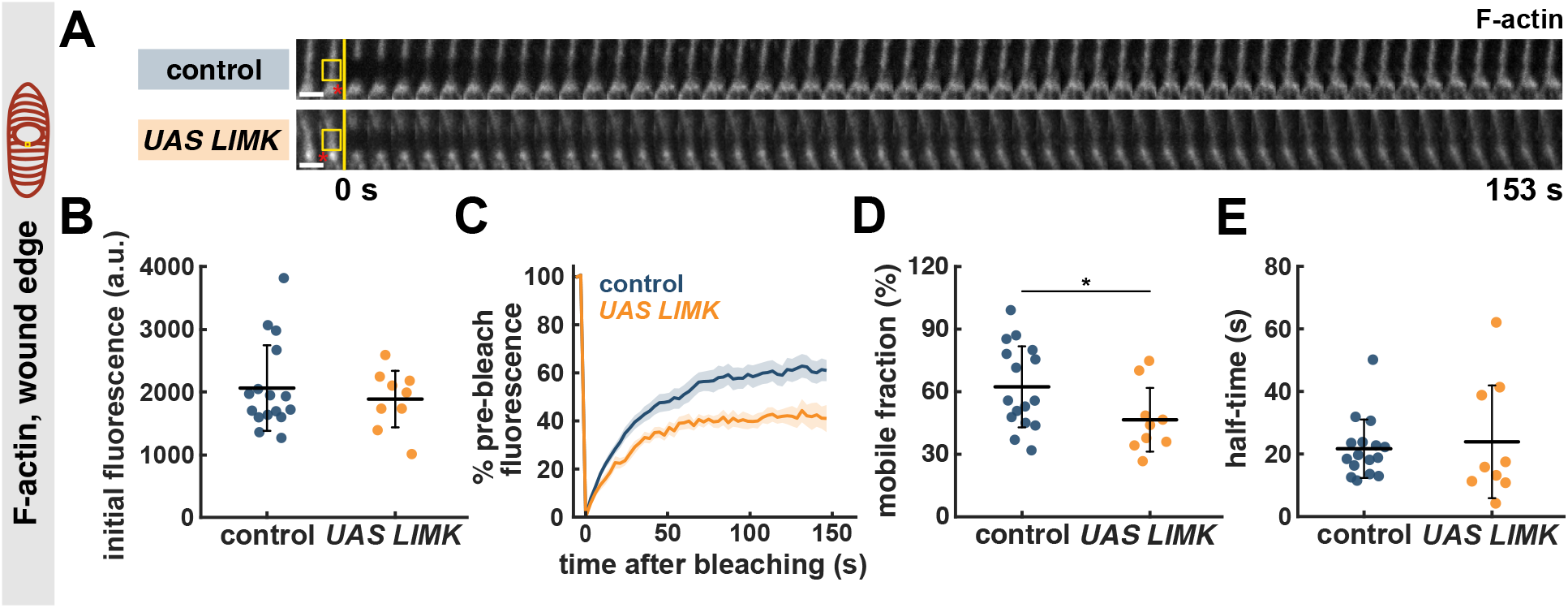
Cofilin activity promotes F-actin turnover at the wound edge. **(A)** Kymographs displaying GFP:UtrophinABD FRAP in a segment of the cable at the wound edge in control (top) or LIMK-overexpressing (bottom) embryos. Red asterisks indicate the position of the wound. Yellow boxes show the photobleached region, yellow lines indicate the time of photobleaching. Anterior left, ventral down. Bars, 3 s. **(B-E)** Initial fluorescence before photobleaching (B), percent of fluorescence recovered over time (C), mobile fraction (D), and half-time of fluorescence recovery (E) for FRAP experiments in control (blue, *n* = 15 wounds) or LIMK-overexpressing embryos (orange, *n* = 9). (B, D-E) Error bars, SD; line, mean. * *P* < 0.05 (Mann-Whitney test). (C) Error bars, SEM.

Changes to tissue mechanics can disrupt wound healing responses (Tetley et al., 2019). To establish if the effect of Cofilin on actin turnover and rapid wound repair is specific to the wound edge, we photobleached GFP:UtrophinABD in epidermal cell-cell junctions of embryos that had not been wounded (Fig. S5A). We found that there were no significant differences in the degree or rate of fluorescence recovery in the intact epidermis of embryos with reduced Cofilin activity when compared to controls (Fig. S5B-E), consistent with the reduced levels of Cofilin at cell-cell junctions (Fig. 1A). These results suggest that Cofilin acts specifically at the wound edge during embryonic wound closure. We used laser ablation of cell-cell junctions to quantify the mechanical properties of the intact epidermis (Fig. S6A). We ablated single cell-cell junctions and measured the velocity of retraction of the ends of the severed junction, which is proportional to the tension released (Hutson et al., 2003; Zulueta-Coarasa & Fernandez-Gonzalez, 2015). We did not find any effects of reducing Cofilin activity on tension levels (Fig. S6B). To establish if the reduction in Cofilin activity affects the viscoelastic properties of the epidermis, we modelled the results of the laser ablation experiments as the damped recoil of an elastic fibre using a Kelvin-Voigt mechanical equivalent circuit (Zulueta-Coarasa & Fernandez-Gonzalez, 2015). Using the model, we measured the asymptotic recoil distance, proportional to the tension-to-elasticity ratio, and a relaxation time proportional to the viscosity-to-elasticity ratio. There were no significant differences in either metric between control embryos and embryos with reduced Cofilin activity (Fig. S6C-D). Together, our data show that Cofilin acts specifically at the wound edge to promote F-actin turnover and rapid wound healing, with no significant effect on the tissue-scale mechanics of the embryonic epidermis.

### Cofilin controls myosin turnover but not tension at the wound edge

F-actin and myosin largely overlap at the wound edge (Kobb et al., 2019). To determine if reducing Cofilin activity disrupts myosin dynamics during wound closure, we quantified myosin:GFP FRAP at the wound edge in controls and in embryos with reduced Cofilin activity (Fig. S3E). We found that the mobile fraction of myosin decreased by 31% when Cofilin activity decreased (44 ± 13% *vs*. 64 ± 15% in controls, *P* = 0.03, Fig. S3F-H), with no significant effect on the half-time of fluorescent recovery (Fig. S3I). Thus, our results show that Cofilin promotes myosin turnover during embryonic wound closure.

Myosin stabilization is associated with increased tension at the wound edge (Kobb et al., 2017). To test if tension at the wound edge increases when Cofilin activity decreases, we used laser ablation to quantify the mechanical properties of the wound margin (Fig. S6E). We found no significant difference in the retraction velocity after ablation in embryos with reduced Cofilin activity with respect to controls (Fig. S6F). Mechanical modelling further suggested that reducing Cofilin activity did not affect tension or viscoelastic properties around the wound (Fig. S6G-H). Together, these data indicate that Cofilin activity does not control force generation or viscoelasticity at the wound edge.

### F-actin turnover must be precisely tuned for rapid wound healing

To determine if precise control of F-actin turnover is key for rapid wound repair, we tested whether increasing F-actin turnover rescued the wound healing phenotype in embryos with reduced Cofilin activity. To increase F-actin turnover, we treated embryos with Latrunculin A (LatA), a toxin that sequesters actin monomers and promotes actin filament severing and depolymerization at low concentrations (Fujiwara et al., 2018). To determine if LatA treatment increased F-actin turnover, we conducted fluorescence recovery after photobleaching (FRAP) experiments in embryos expressing GFP:UtrophinABD and treated with 50% DMSO (controls) or with 750 μM of LatA in 50% DMSO (Fig. S7A). In FRAP experiments, embryos treated with LatA displayed an F-actin mobile fraction 48% higher than DMSO controls (54 ± 15% *vs*. 37 ± 6%, respectively, *P* = 0.01, Figure S7B-D), with no significant effects on the half-time of fluorescence recovery (Fig. S7E). Thus, LatA treatment is an effective method to destabilize F-actin. Increasing F-actin turnover was associated with a 67% reduction in the rate of wound closure (*P* = 0.007, Fig. 4A-B, E-F, Video S2). Consistent with our previous findings, reducing Cofilin activity slowed down wound closure by 37% (*P* = 0.02, Fig. 4A, C, E-F, Video S2). Thus, both excessive and insufficient F-actin turnover are detrimental for rapid wound repair. Strikingly, increasing F-actin turnover with LatA in embryos with reduced Cofilin activity rescued the rate of wound closure to control levels (Fig. 4A-F, Video S2). Overall, our results suggest that there is an optimal degree of F-actin turnover that is necessary for rapid wound repair.

**Figure 4.**
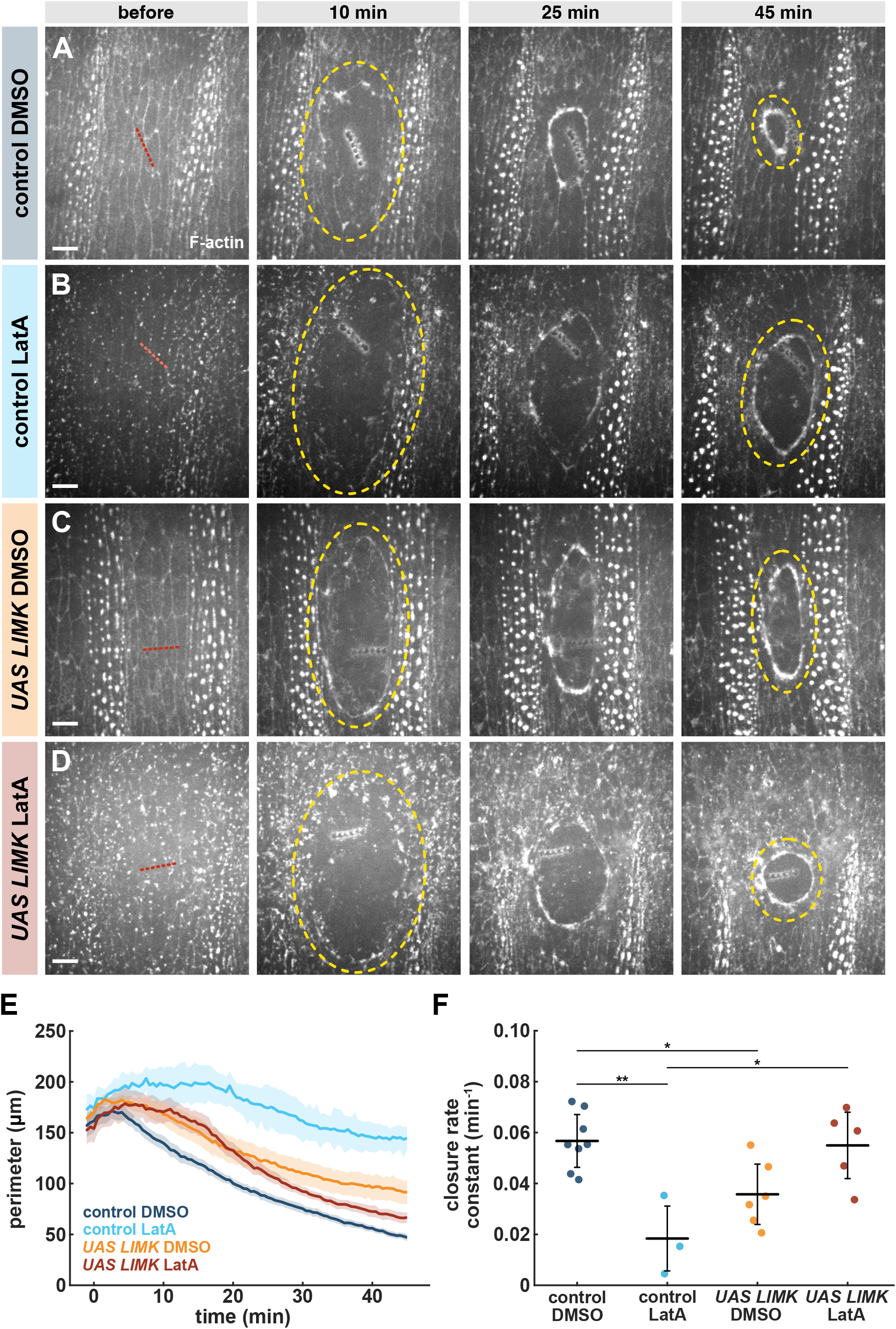
Precise levels of F-actin turnover are necessary for rapid wound healing. **(A-D)** Wound closure in control (A-B) or LIMK-overexpressing embryos (C-D) injected with 50% DMSO (A, C) or 750 μM LatA (B, D), and expressing GFP:UtrophinABD. Red dotted lines indicate wound sites, yellow dashes outline the wounds. Time is with respect to wounding. Anterior left, ventral down. Bars, 10 μm. **(E-F)** Wound perimeter over time (E) and wound closure rate constant (F), for control embryos treated with DMSO (blue, *n* = 8 wounds) or LatA (cyan, *n* = 3), and for LIMK-overexpressing embryos treated with DMSO (orange, *n* = 6) or with LatA (red, *n* = 5). (E) Error bars, SEM. (F) Error bars, SD; line, mean. * *P* < 0.05, ** *P* < 0.01 (Dunn’s test).

### Computational modelling suggests that Cofilin may control F-actin rigidity to drive rapid wound healing

Cofilin and F-actin turnover could contribute to wound healing through effects on actin filament length (Jermyn et al., 2020; Kremneva et al., 2014; Robaszkiewicz et al., 2020), disassembly rate (Carlier et al., 1997; Wioland et al., 2017), filament severing (Pavlov et al., 2007), or F-actin rigidity (Brannon et al., 2011; McCullough et al., 2008). To investigate how different actin-associated parameters may affect embryonic wound healing, we developed a computational model of wound repair with molecular resolution. We implemented our model using Cytosim (Nedelec & Foethke, 2007), which allows the simulation of actin networks containing filaments of tunable length, (dis)assembly rate and rigidity, and includes motor proteins, actin crosslinkers, and actin-severing proteins. Using Cytosim, we created an elliptical network of actin filaments to simulate the wound healing response (see Methods and Table S1). The *Drosophila* embryonic epidermis is anisotropic, with F-actin and myosin polarized parallel to the dorsal-ventral axis of the embryo (Simone & DiNardo, 2010). As a consequence, epidermal wounds in the *Drosophila* embryo are elongated along the dorsal-ventral axis (Abreu-Blanco et al., 2012; Fernandez-Gonzalez & Zallen, 2013). To simulate the mechanical anisotropy of the epidermis, we generated actin networks in Cytosim with an initial circularity comparable to that of a wound *in vivo* (see Methods). Parameter values were extracted from the literature (Table S1), with a final calibration step to obtain closure times consistent with our *in vivo* experiments. To examine how different parameters affected the contraction of the actin filament network, we individually varied initial filament length, actin filament disassembly rate, filament severing, and filament rigidity. Specifically, we investigated how potential effects of reducing Cofilin activity (increasing initial filament length, reducing the rate of actin filament disassembly, reducing actin severing or increasing filament rigidity) would affect the rate of network contraction (Figs. 5A-D and S8A-D). The model predicted that reducing actin severing or increasing filament rigidity would both be detrimental for actomyosin network contraction (Fig. 5E-F, H-I, Videos S3 and S4). In contrast, increasing the length of actin filaments or reducing the rate of F-actin disassembly did not negatively impact the contraction of the network (Fig. S8A-J). Importantly, the model predicted that an increase in filament rigidity, but not a reduction of actin severing, would alter the shape of the network throughout its contraction, with flexible networks significantly more circular than rigid ones (*P* < 0.01, Fig. 5G, J). Further increasing network rigidity led to a greater reduction in wound circularity and the rate of wound closure (Fig. S9A-E). Thus, our model predicts that Cofilin may control the rate of wound closure by inducing actin severing, with no effect on wound shape; or by promoting a flexible F-actin network and facilitating wound rounding as the wound closes.

**Figure 5.**
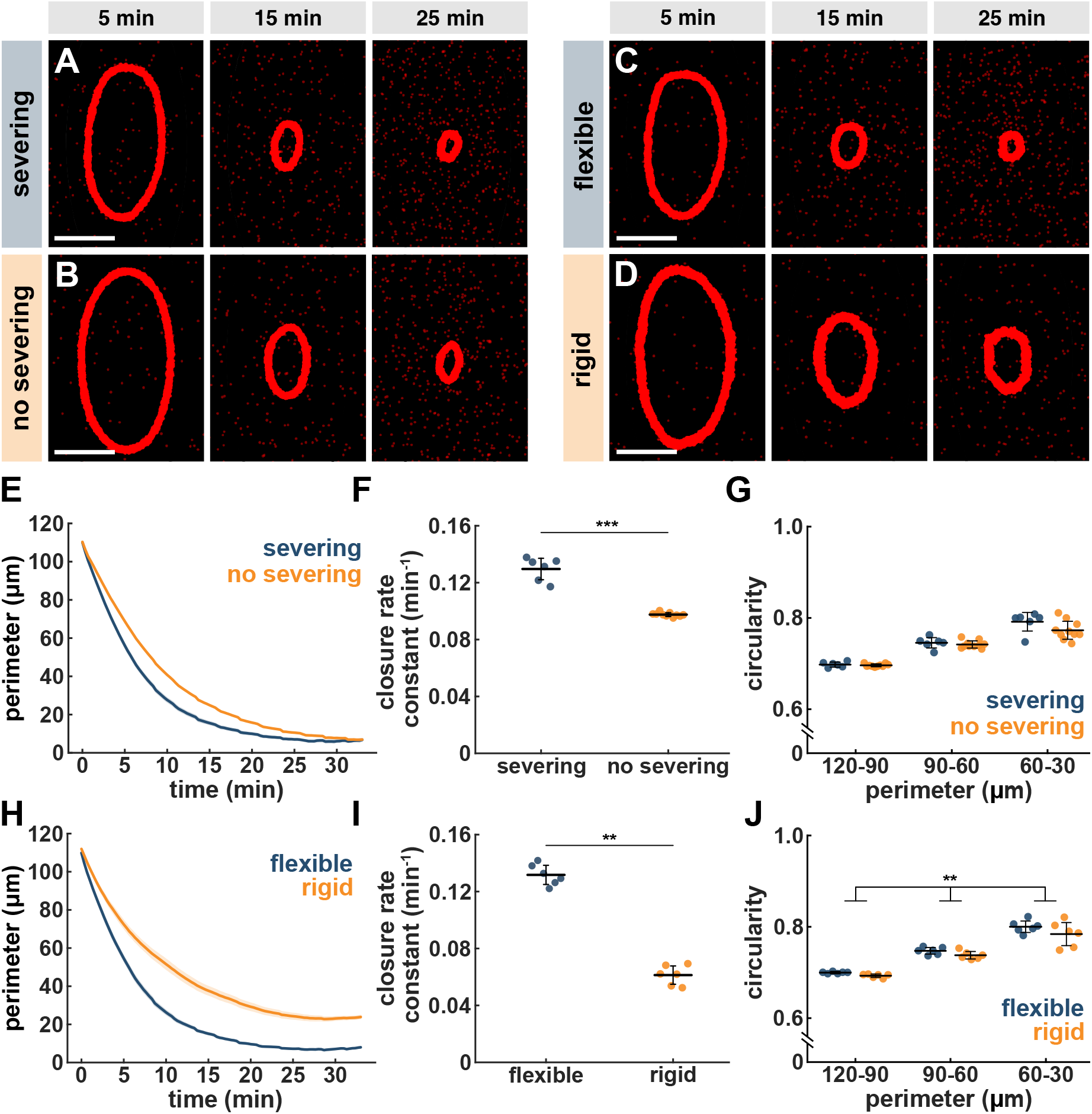
Computational modelling predicts that Cofilin induces F-actin severing or controls F-actin rigidity to drive rapid wound repair. **(A-D)** Wound closure in simulations with (A) or without (B) F-actin severing, or with flexible (C) or rigid (D) F-actin. Red represents an actin-binding protein to visualize network contraction. Time is with respect to the onset of wound healing. Bars, 10 μm. **(E-J)** Wound perimeter over time (E, H), wound closure rate constant (F, I), and simulation averages of wound circularity binned by wound perimeter (G, J) for simulations with (blue, *n* = 6 simulations) or without (orange, *n* = 10) actin severing (E-G), and with flexible (blue, *n* = 6) or rigid (orange, *n* = 6) F-actin (H-J). (E, H) Error bars, SEM. (F-G, I-J) Error bars, SD; line, mean. ** *P* < 0.01, *** *P* < 0.001 (Mann-Whitney (F, I) or Fisher’s method combining *P*-values for all 3 bins (J)).

### Cofilin facilitates wound rounding associated with rapid repair

To distinguish between potential effects of Cofilin on wound healing by affecting F-actin severing or rigidity, we measured the circularity of wounds in living embryos with reduced Cofilin activity (Fig. 6A-B). Reducing Cofilin activity was associated with a 10-19% reduction in circularity when comparing wounds at specific time points during wound closure (*P* < 0.005, Fig. 6C). Similarly, when comparing wounds of similar perimeter, reducing Cofilin activity resulted in wounds that were 8-12% less circular than in controls (*P* < 0.01, Fig. 6D). We obtained similar results when we examined wound morphology in *tsr* ^*+/-*^ mutants (Fig. S10A-D). These results are consistent with our model predictions if Cofilin limits F-actin rigidity at the wound edge (Fig. 5J). The circularity defects when Cofilin activity or levels decrease *in vivo* may be exacerbated by the dorsal-ventral pre-stress that morphogenetic movements on the dorsal side of the embryo impose on the epidermis (Young et al., 1993). Overall, our data suggest that Cofilin may contribute to rapid wound closure by maintaining a flexible F-actin network during embryonic wound healing. Notably, we found that increasing F-actin turnover with LatA treatment rescued wound circularity in embryos with reduced Cofilin activity (Fig. S11A-E), suggesting a potential link between the effects of Cofilin on F-actin turnover and rigidity.

**Figure 6.**
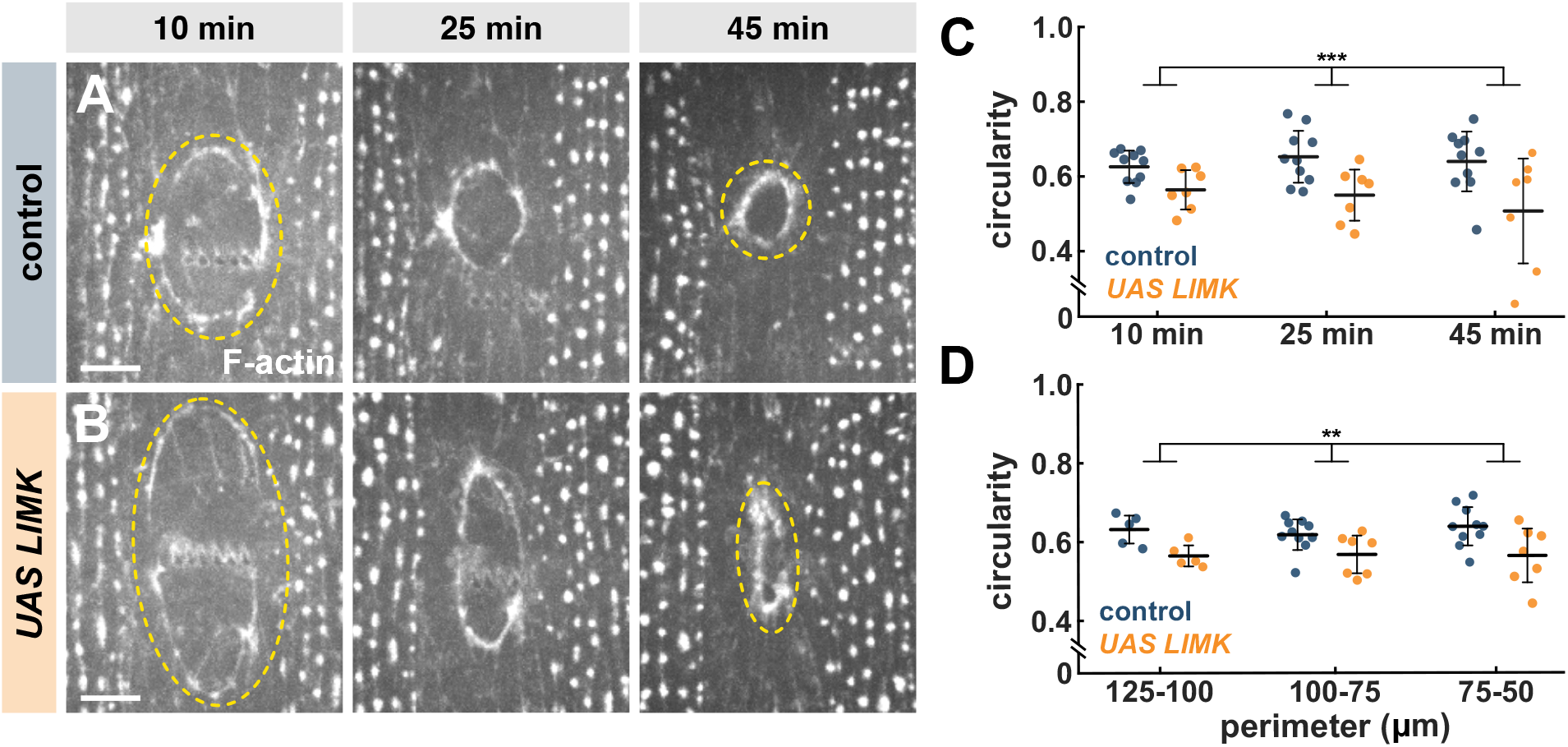
Cofilin activity facilitates wound rounding. **(A-B)** Wounds at specific times during tissue repair in control (A) and LIMK-overexpressing (B) embryos. Embryos express GFP:UtrophinABD. Yellow dashes outline the wounds. Anterior left, ventral down. Bars, 10 μm. **(C-D)** Wound circularity at specific times during tissue repair (C), or for wounds of certain perimeter (D) in control (blue, *n* = 10 wounds) or LIMK-overexpressing embryos (orange, *n* = 7). Error bars, SD; line, mean. ** *P* < 0.01, *** *P* < 0.005 (Fisher’s method combining *P*-values for all 3 bins).

### Cofilin limits F-actin rigidity *in vivo*

Our results suggest that Cofilin promotes F-actin turnover to limit F-actin rigidity and facilitate rapid embryonic wound healing. To establish if reducing Cofilin activity affects F-actin rigidity *in vivo*, we measured the periodicity of F-actin fluorescence fluctuations at the wound edge (Bohdan et al., 2016; Sun et al., 2024; Zhang et al., 2017). Flexible actin filaments display greater (and thus less frequent) thermal fluctuations than rigid filaments (McCullough et al., 2008) (Fig. 7A). Thus, if actin filaments were fluorescently labeled, the period of fluorescence fluctuations in a diffraction-limited spot should be greater for flexible F-actin networks than for rigid ones (Fig. 7A′). We quantified GFP:UtrophinABD fluctuations at the wound edge in control embryos and in embryos overexpressing LIMK using a spinning disk confocal microscope (Needleman et al., 2009; Sisan et al., 2006). We selected wounds at 50% closure and acquired an image of the F-actin cable around the wound. After stopping the spinning disk, we selected pinholes overlaying the F-actin cable (Fig 7B). For each pinhole, we continuously collected 1600 images with 5 msec exposures (Fig. 7B, cutout). The amplitude of F-actin fluorescence fluctuations was dramatically reduced in fixed embryos with respect to living ones (Fig. 7C). To quantify the F-actin fluctuation period, we measured the autocorrelation of the fluorescence fluctuations (Fig. 7D). We found that the autocorrelation decay time was 23% lower in embryos with reduced Cofilin activity with respect to controls (*P* = 0.02, Fig. 7-E), consistent with a reduction in fluctuation period and an increase in F-actin rigidity when Cofilin activity decreased. Similarly, we observed a 20% reduction in autocorrelation decay time in *tsr* ^*+/-*^ embryos when compared to controls (*P* = 0.04, Fig S12A-B). Together, our results indicate that Cofilin maintains a flexible F-actin network at the wound edge to facilitate rapid wound repair.

**Figure 7.**
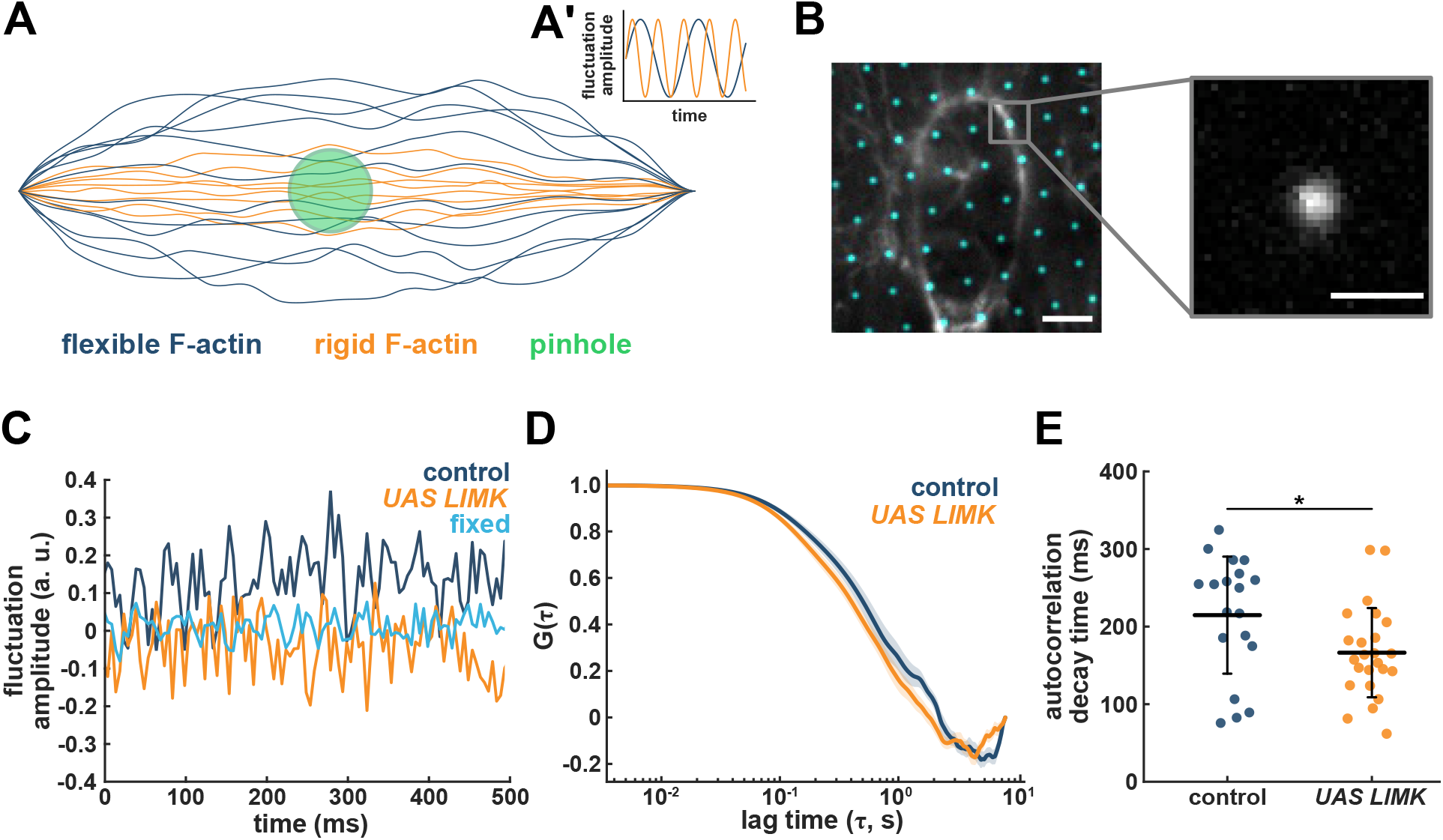
Cofilin controls F-actin rigidity *in vivo*. **(A)** Cartoon depicting the impact of F-actin rigidity on thermal fluctuations (A) and fluorescence fluctuations in a diffraction-limited spot (A’). **(B)** F-actin cable around a wound in an embryo expressing GFP:UtrophinABD and imaged while the disk in a spinning disk confocal rotated (grayscale); and arrested disk pinholes (cyan), with sample signal collected from a pinhole (right). Anterior left, ventral down. Bars, 10 μm (left) and 1 μm (right). **(C)** Representative sample fluorescence fluctuations measured through an arrested spinning disk pinhole in a fixed embryo (cyan), or in living control (blue) and LIMK-overexpressing (orange) embryos expressing GFP:UtrophinABD. **(D-E)** Mean autocorrelation, *G* (D) and autocorrelation decay times (E) for GFP:UtrophinABD fluctuations at the wound edge in control (blue, *n* = 19 pinholes in 13 wounds) and LIMK-overexpressing embryos (orange, *n* = 24 pinholes in 18 wounds). (D) Error bars, SEM. (E) Error bars, SD; line, mean. * *P* < 0.05 (Mann-Whitney test).

## Discussion

Here we show that Cofilin contributes to rapid wound repair by promoting F-actin turnover and flexibility specifically at the wound edge. The mechanisms that polarize Cofilin to the wound margin remain unclear. F-actin polarization to the wound edge depends on calcium (Hunter et al., 2015; Razzell et al., 2018), reactive oxygen species (Hunter et al., 2018), and signalling by the small GTPases Rap1 (Rothenberg et al., 2023) and Rho1 (Abreu-Blanco et al., 2012; Wood et al., 2002). Cofilin is a small protein that is able to freely diffuse (Bamburg, 1999). Thus, the affinity of Cofilin for F-actin could localize Cofilin at the wound margin. Alternatively, localized activation could mediate the polarization of Cofilin. Cofilin activation is mediated by the phosphatase Slingshot (Niwa et al., 2002), and Slingshot activity can be controlled by calcium (Wang et al., 2005), reactive oxygen species (Kim et al., 2009), or F-actin binding (Niwa et al., 2002). However, our data indicate that in the embryonic epidermis Cofilin is largely active prior to wounding, suggesting that controlling localization, rather than activity, is key for the specific role that Cofilin plays at the wound margin.

Our results demonstrate that Cofilin also controls myosin turnover around the wound. We previously showed that myosin turnover is regulated by tension, with increasing tension slowing down myosin turnover (Fernandez-Gonzalez et al., 2009; Kobb et al., 2017). This is possibly due to an increase in the myosin duty ratio (the fraction of time that the motor is bound to the filament) under high tension as a consequence of the slower release of ADP from the motor head (Kovács et al., 2007). Thus, our results suggest that at least two mechanisms control the dynamics of the actomyosin cytoskeleton around embryonic wounds: (1) Cofilin mobilizes actin and myosin at the wound margin; and (2) tension stabilizes myosin motors. Cofilin-based mobilization may allow myosin motors or actin crosslinkers to dissociate from and reassociate with actin filaments (Mendes Pinto et al., 2012), thus inducing productive contraction. This would be balanced by tension-based stabilization of motors and crosslinkers to ratchet contraction (Zulueta-Coarasa & Fernandez-Gonzalez, 2018). Whether myosin turnover and stabilization are coordinated is not understood. Cofilin binds and severs actin filaments in a tension-sensitive manner, with preferential binding and faster severing of relaxed filaments (Hayakawa et al., 2011). Tension at the wound edge increases as wound healing progresses (Kobb et al., 2017), suggesting that Cofilin-based actin severing may slow down as the wound closes, consistent with the slower turnover of myosin at later stages of tissue repair. Further studies investigating whether tension regulates Cofilin dynamics at the wound edge will shed light on the mechanisms that balance cytoskeletal turnover and stability to drive rapid wound repair.

We demonstrate that Cofilin maintains a flexible F-actin network around the wound. The flexibility of the F-actin network may facilitate filament packing as myosin motors slide the filaments past each other. Additionally, flexible actin filaments may oppose lower resistance to myosin- or crosslinker-based filament alignment (Butt et al., 2010; Lieleg et al., 2007), a process in which Cofilin has been implicated (Mseka et al., 2007) and that is associated with force generation (Bidone et al., 2014; Spira et al., 2017; Stachowiak et al., 2014). Super-resolution analysis of the actin network architecture (Garlick et al., 2022; Hui et al., 2023) at the wound edge will reveal whether Cofilin promotes filament alignment and compaction to facilitate rapid embryonic wound healing.

An outstanding question is whether the effects of Cofilin on F-actin turnover and flexibility during wound repair are related. The architecture and organization of F-actin networks can affect their contraction (Koenderink & Paluch, 2018; M. Murrell et al., 2015) and crosslinking proteins can modulate force transmission throughout a network (M. P. Murrell & Gardel, 2012). The effect of crosslinkers on F-actin networks is influenced by filament density and length, with longer filaments being more heavily crosslinked (Kasza et al., 2010). The presence of crosslinkers also increases the rigidity of F-actin networks *in vitro* (Gardel et al., 2004; Schmoller et al., 2009) and in cells (Katsuta et al., 2023), and reduces turnover (Katsuta et al., 2023; Tilney et al., 2003). Thus, by changing filament conformation (Brannon et al., 2011; McCullough et al., 2008) or dynamics in a way that limits crosslinking (Hylton et al., 2022), Cofilin could maintain a flexible F-actin network, facilitating contraction and packing. Additionally, a high crosslinker density has been associated with reduced contractility (Bendix et al., 2008; Ennomani et al., 2016), likely because crosslinkers hinder F-actin bundle bending (Akenuwa & Abel, 2023), which is an important symmetry breaking mechanism that contributes to actomyosin contraction both *in vitro* (M. P. Murrell & Gardel, 2012) and *in vivo* (Costa et al., 2002). Investigating how different crosslinkers contribute to the wound healing response (Lehne & Bogdan, 2023), and establishing the interplay between crosslinkers and Cofilin at the wound edge, will contribute to our understanding of how the material properties of the cytoskeleton are established and regulated *in vivo*, and how they impact tissue morphogenesis.

## Methods

### Fly stocks

Flies were maintained at room temperature and embryos were collected on apple juice agar plates kept overnight at room temperature. For live imaging, we used *sqh-GFP:utrophinABD* (Rauzi et al., 2010), *sqh-sqh:GFP* (Royou et al., 2004), *endo-E-cadherin:tdTomato* (Huang et al., 2009), *endo-cofilin:GFP* (Ikawa & Sugimura, 2018; Morin et al., 2001) and *endo-aip1:GFP* (Bloomington *Drosophila* Stock Center #50824) (Buszczak et al., 2007). For LIMK overexpression, we used *UAS-LIMK* (Bloomington *Drosophila* Stock Center #9116) (Ng & Luo, 2004). *UAS-LIMK* was ubiquitously driven with *daughterless-Gal4* (Wodarz et al., 1995). We used FlyBase to find information on stocks and gene expression (Öztürk-Çolak et al., 2024).

### Embryo mounting and injections

*Drosophila* embryos at stage 14-15 were dechorionated in 50% bleach for 2 minutes and thoroughly rinsed. Embryos were glued ventral-lateral side down to a glass coverslip using heptane glue and covered with a 1:1 mix of halocarbon oil 27 and 700 (Sigma-Aldrich). For injections, embryos were dehydrated for 8-9 minutes inside a plastic container with silica (Drierite). After dehydration, embryos were covered with the same 1:1 mix of halocarbon oil. Injections were done with a PV820 injector (World Precision Instruments) coupled to a spinning disk confocal microscope. Embryos were injected in the perivitelline space with 750 μM of Latrunculin A (Tocris Bioscience) dissolved in 50% DMSO, or with 50% DMSO as a control. Drugs are predicted to be diluted by 50-fold in the perivitelline fluid (Foe & Alberts, 1983). All injections were followed by a 10-minute incubation period at room temperature before imaging.

### Time-lapse imaging

Embryos were imaged at room temperature using a Revolution XD spinning disk confocal microscope (Andor Technology) with a CSU-X1 confocal head and a 60X oil immersion lens (Olympus, NA 1.35). The microscope was controlled with Metamorph (Molecular Devices). Sixteen-bit Z-stacks were acquired at 0.5 μm steps every 3, 4 or 30 seconds (11 or 21 slices per stack). Maximum intensity projections were used for analysis.

### Laser ablation

Laser ablations were done using a pulsed Micropoint nitrogen laser (Andor Technology) tuned to 365 nm. For wounding, 10 pulses were delivered at six spots along a 13 μm line. For spot laser ablations, 10 pulses were delivered at a single point. Samples were imaged immediately before and after ablation.

### FRAP

Photobleaching experiments were done using a FRAPPA system (Andor) and a 488 nm laser. A 2.7 × 2.7 μm^2^ region (10 x 10 pixels) was photobleached using a dwell time of 500 μs/pixel. Two Z-stacks were acquired 3 s apart prior to photobleaching. Photobleached regions were imaged immediately after photobleaching and every 3 s thereafter for at least 2 min.

### F-actin fluctuation measurements

Living embryos expressing GFP:UtrophinABD were wounded and imaged in a spinning disk confocal microscope until wound area reached 50% of its maximum value. A single Z-slice image was collected using a 100X oil-immersion lens (NA 1.4, Olympus) to map the position of the wound edge. Disk rotation was stopped, and 1600 single Z-slice images were captured sequentially, with an exposure time of 5 ms. The pinholes that overlapped the wound edge were identified by overlaying the first image of the sequence with the image acquired before the disk was stopped.

### Embryo fixation and immunofluorescent staining

Embryos at stage 14-15 were dechorionated as described above and fixed for 40 minutes in a 1:1 mix of heptane and 37% formaldehyde in phosphate buffer. Embryos were devitellinized manually with the aid of a glass needle, stained with primary antibodies overnight at 4ºC, and incubated with secondary antibodies for 1 hour at room temperature. Primary antibodies were rabbit anti-Dm-Cofilin (Figard et al., 2019) (1:500, a gift from Anna Sokac), and mouse anti-Pi-Hs-Cofilin1 (1:100, sc-271921, Santa Cruz Biotechnology). Fluctuation analysis in fixed embryos was conducted in embryos stained with Phalloidin 488 (1:500, Invitrogen). Embryos were mounted in ProLong Gold (Molecular Probes) between two coverslips. Sixteen-bit Z-stacks were acquired at 0.2 μm steps (27 slices per stack). Single Z-slices were used for quantitative analysis.

### Quantitative image analysis

Image analysis was done using PyJAMAS, an open-source image analysis platform that we develop (Fernandez-Gonzalez et al., 2022). To quantify the rate of wound closure and fluorescence intensities around the wound, we generated maximum intensity projections of the stacks and traced the wound edge in each frame of the time-lapse sequences using the semi-automated LiveWire tool (Fernandez-Gonzalez & Zallen, 2011), integrated in PyJAMAS. The LiveWire method uses Dijkstra’s optimal path search algorithm (Dijkstra, 1959) to find the brightest pixel path connecting any two pixels, and can be used to rapidly trace the wound edge (Scepanovic et al., 2021). We measured the rate of wound closure by fitting an exponential decay function to the curve that quantifies wound area over time and extracting the decay constant (Hunter et al., 2018). We measured the circularity, *C*, of wounds as:

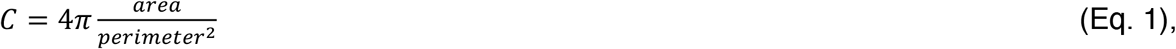

with a perfect circle having a circularity of 1, and shapes deviating from circles having lower circularity values. Fluorescence at the wound margin was calculated as the mean pixel value under a 0.6 μm-wide mask generated by the LiveWire algorithm. Intensity values were background-subtracted using the image mode, and corrected for photobleaching by dividing by the mean image intensity at each time point. For the quantification of Cofilin and AIP1 accumulation at the wound edge, a small region outside of the embryo was used for background correction calculations.

To measure retraction velocity after laser ablation, we tracked the position of the two tricellular junctions adjacent to the ablation site. Additionally, we modelled junctions as viscoelastic elements using a Kelvin-Voigt model (Zulueta-Coarasa & Fernandez-Gonzalez, 2015), which describes the distance *L* between the ends of the severed junction at time *t* after ablation as:

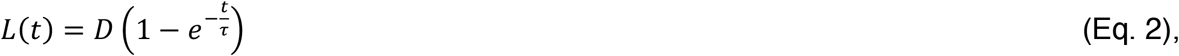

where *D* is the asymptotic distance of retraction, proportional to the tension sustained by the junction, and *τ* is a relaxation time given by the viscosity-to-elasticity ratio.

For FRAP analysis, the fluorescence intensity in the photobleached region at time *t, f(t)*, was measured as:

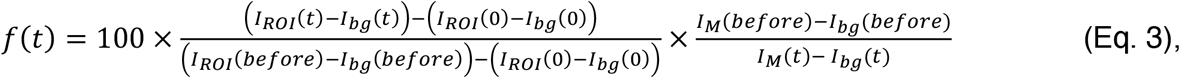

where *t=before* is the time immediately before photobleaching, *t* = 0 is the time immediately after photobleaching, *I*_*ROI*_ is the mean pixel value within the photobleached region, *I*_*bg*_ is the background signal, calculated as the mean pixel value within a 10×10-pixel region outside the embryo, and *I*_*M*_ is the mean image intensity. The mobile fraction (*M*) was obtained from fitting the fluorescence recovery curve with an exponential:

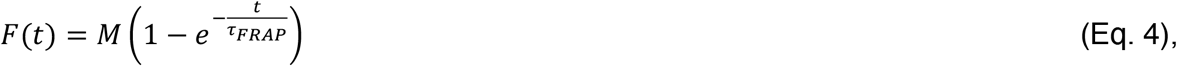

where *τ*_*FRAP*_ is a characteristic time scale of fluorescence recovery. The half-time of fluorescence recovery, *t*_*1/2*_, was calculated as (Pines et al., 2012):

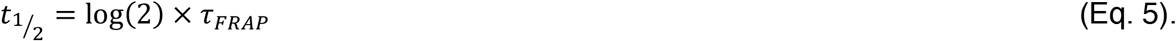

For F-actin fluctuation analysis, fluorescence intensities imaged through wound edge pinholes were measured using PyJAMAS. Raw intensities for each pinhole were background-subtracted using the image mode to account for camera noise. Photobleaching normalization was conducted by dividing the fluctuation value at time *t* by the mean of all the wound edge pinhole intensities at that specific time point. Fluctuations for each pinhole trace were calculated by subtracting the temporal mean from the background-corrected and photobleaching normalized intensities. Autocorrelation results were normalized to a maximum value of 1, and were fitted to the model:

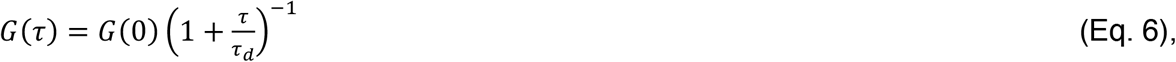

where *G*(0)=1, and *τ*_d_ is the autocorrelation decay time, related to the period of fluctuation.

### Computational modelling

Contraction of the actomyosin cable at the wound edge was modelled with Cytosim, an open-source platform to model cytoskeletal dynamics (Lugo et al., 2023; Nedelec & Foethke, 2007). The models created in Cytosim are agent-based models, which allow for the study of interactions between autonomous agents. Cytosim provides “functional units”, including actin filaments, myosin motors, crosslinkers, and actin-severing proteins. The functional units in Cytosim are instantiated to create individual agents. Agents possess certain attributes (*e*.*g*. for an actin filament, an initial length, rates of assembly and disassembly, a rigidity, etc., Table S1), whose values contribute to the overall energy of the system. Cytosim reconstructs the dynamics of the different agents by minimizing the overall energy of the system. Control parameter values were adjusted in calibration experiments to approach the *in vivo* rates of network contraction. For each condition, we simulated actin bundles made up of 8 filaments. The initial actin levels were the same in all simulations, consistent with our *in vivo* measurements (Figs. 2E-H and 3B). Thus, for instance, when we simulated scenarios with actin filament length halved, we doubled the number of bundles to preserve the same amount of actin. Similarly, actin filaments were arranged into an ellipse with dimensions typical of our *in vivo* measurements: a major axis of 50 μm and a minor axis of 20 μm, for a circularity of 0.75 (Fig. 6C). Myosin motors were assembled into myosin minifilaments of 8 motors each. Actin bundles and myosin minifilaments were distributed randomly along the ellipse in each trial, such that at the start of the simulations the actin filaments were tangential to the ellipse. Crosslinkers were distributed randomly within each bundle. Simulations were run with a time step of 0.1 seconds, evolving stochastically according to Langevin dynamics (Nedelec & Foethke, 2007). Frames showing the simulated networks at 20 second intervals were exported and analyzed using PyJAMAS.

### Statistical analysis

To assess the significance of differences between two unpaired groups, we used a non-parametric Mann-Whitney test. For multigroup comparisons, we used a Kruskal-Wallis test to reject the null hypothesis, and post-hoc Dunn’s tests for pairwise comparisons. To compare parameter values over perimeter bins, we used Fisher’s method for P-value combination (Fisher, 1932). Single comparisons were made using the Mann-Whitney test, and the obtained P-values (*P*_*i*_) were combined into a joint test to determine the presence of a global effect. The Fisher method statistic (*X*) follows a χ^2^ distribution with *ν = 2k* degrees of freedom, and was calculated as:

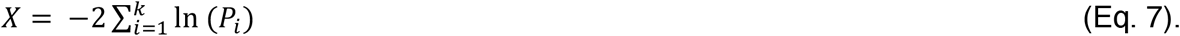

## Acknowledgments

We are grateful to Ray Hawkins for help with image analysis. We thank Negar Balaghi, Veronica Castle, Ray Hawkins, Sasha Korolov, Willow Peterson, and Gordana Scepanovic for comments on the manuscript, and Tony Harris, Shane Hutson, and Alison McGuigan for helpful discussions. AMC was partially supported by a PhD fellowship from the Portuguese Foundation for Science and Technology (FCT, PD/BD/135448/2017). JHS was partially supported by scholarships from the Translational Biology and Engineering Program of the Ted Rogers Centre for Heart Research and the University of Toronto Excellence Award. This work was funded by grants to RFG from the Canadian Institutes of Health Research (156279 and 186188) and the Canada Foundation for Innovation (30279). RFG is the Canada Research Chair in Quantitative Cell Biology and Morphogenesis.

## Notes

### Competing Interest Statement

The authors have declared no competing interest.

### Summary of Updates

We showed that not only a reduction in Cofiilin activity (in embryos overexpressing the kinase that phosphorylates and inactivates Cofilin, LIMK), but also a reduction in Cofilin levels (in *cofilin* heterozygous mutants) slows down wound repair. *cofilin* mutants displayed reduced F-actin turnover specifically at the wound edge. The reduction in F-actin turnover around the wound was associated with an increase in F-actin rigidity, as measured by fluorescence fluctuation spectroscopy. Thus, our results indicate that Cofilin contributes to rapid embryonic wound healing by maintaining a flexible F-actin network at the wound edge that can be efficiently packed as it contracts.

