## Supplementary material for "Cofilin promotes actin turnover and flexibility to drive coordinated cell movements *in vivo*": Figures S1-S12, Table S1, supplementary video legends

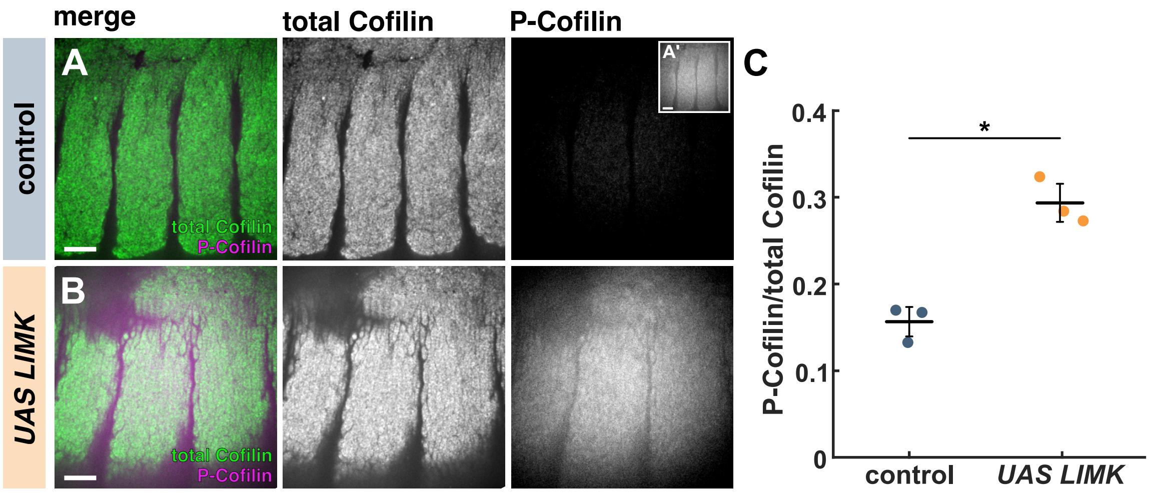


**Figure S1. LIMK overexpression inactivates Cofilin in the *Drosophila* embryonic epidermis. (A-B)** Control (A) and LIMK-overexpressing (B) embryos fixed and stained with antibodies against total Cofilin (green in merge, middle) and P-Cofilin (magenta in merge, right). Inset in A shows a contrast-stretched image of P-Cofilin fluorescence in control embryos. Anterior left, ventral down. Bars, 200 µm. **(C)** P-Cofilin-to-total Cofilin ratio in control ($n$ = 3 embryos) and LIMK-overexpressing embryos ($n$ = 3). Error bars, SD; line, mean. * $P < 0.05$ (Mann-Whitney test).

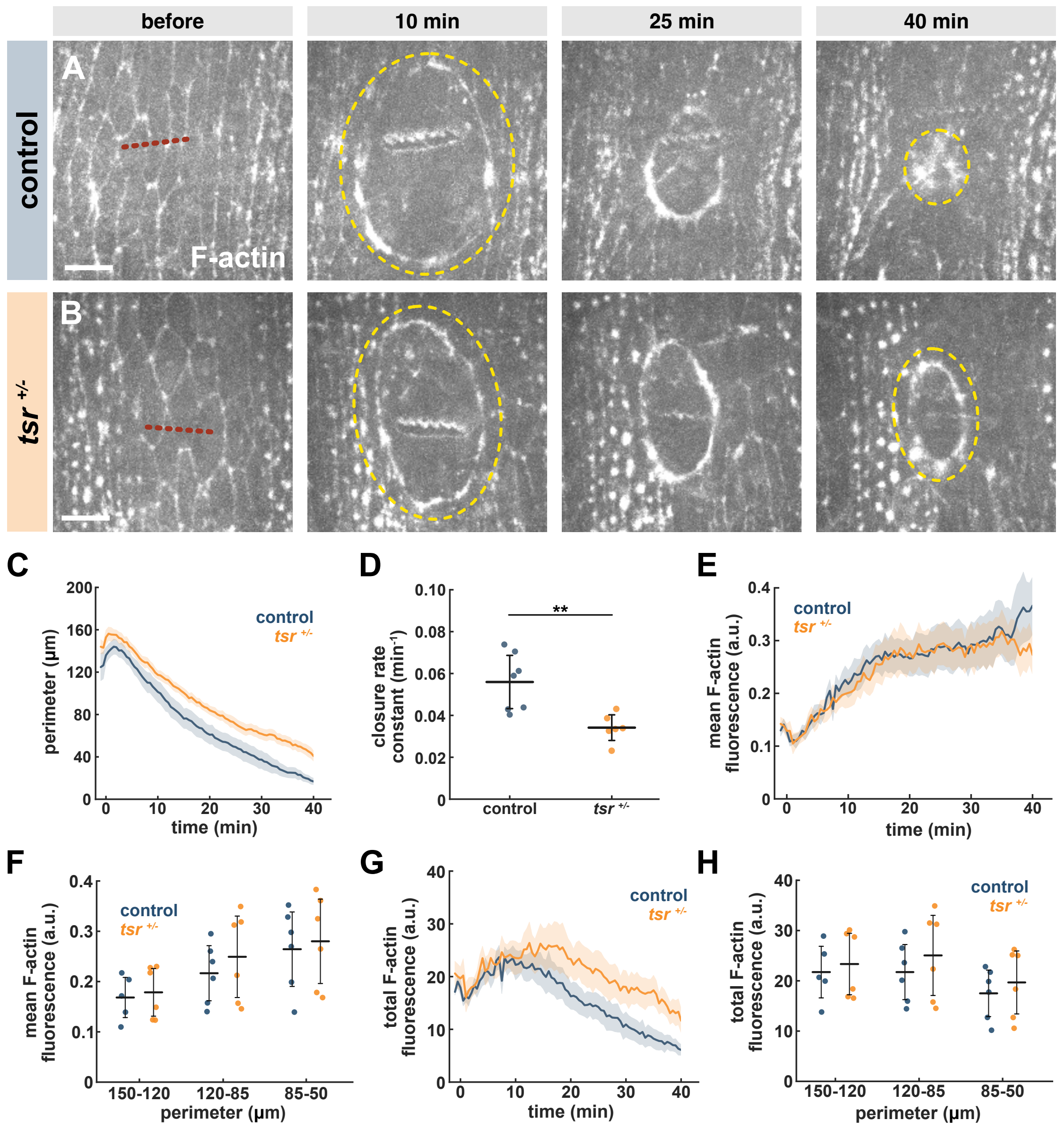


**Figure S2. Cofilin is necessary for rapid wound closure. (A-B)** Wound closure in control (A) and *tsr* $^{+/-}$ mutant (B) embryos expressing GFP:UtrophinABD. Red dotted lines indicate wound sites, yellow dashes outline the wounds. Time is with respect to wounding. Anterior left, ventral down. Bars, 10 µm. **(C-H)** Wound perimeter over time (C), wound closure rate constant (D), mean or total F-actin fluorescence at the wound edge over time (E or G, respectively), or embryo averages binned by perimeter (F or H, respectively) in control (blue, $n = 7$ wounds) or *tsr* $^{+/-}$ mutant embryos (orange, $n = 6$). (C, E, G) Error bars, standard error of the mean (SEM). (D, F, H) Error bars, SD; line, mean. ** $P < 0.01$ (Mann-Whitney test).

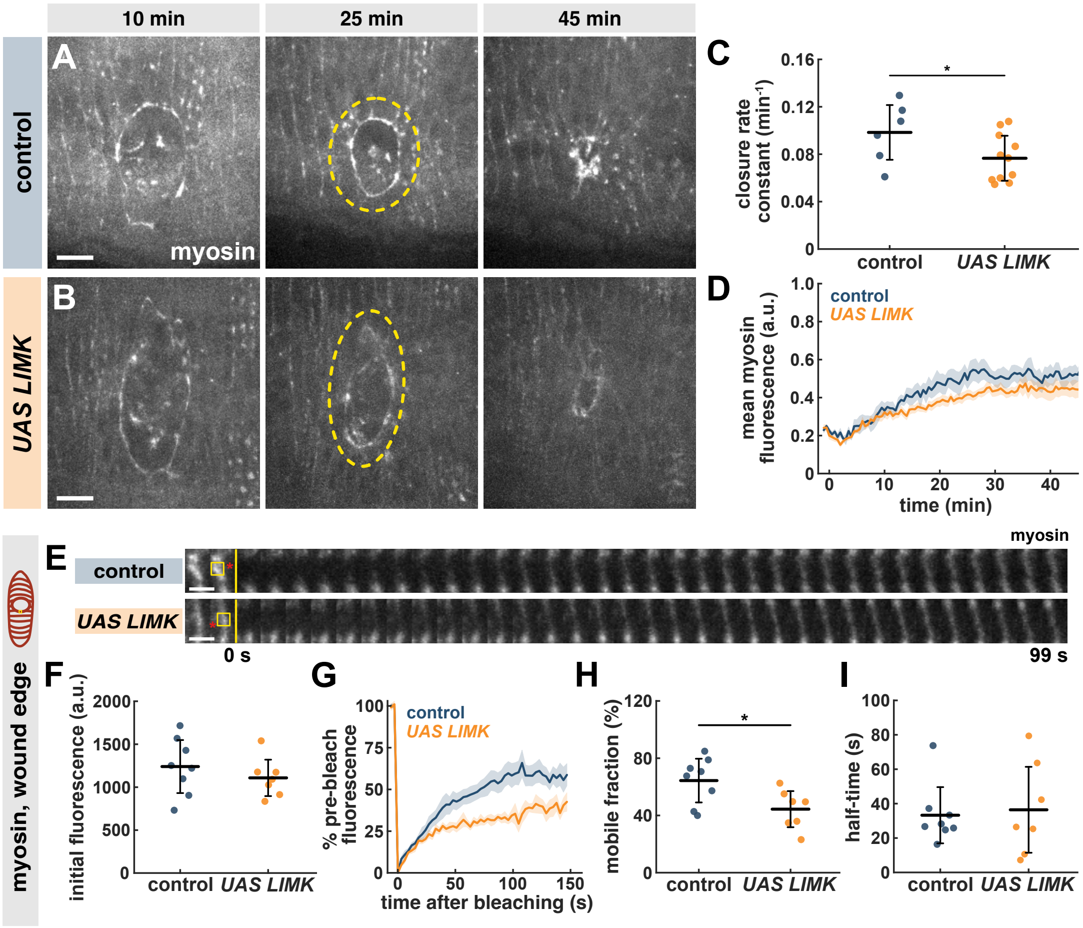


**Figure S3. Cofilin controls myosin turnover but not polarization during embryonic wound closure. (A-B)** Wound closure in control (A) and LIMK-overexpressing (B) embryos expressing a fluorescent myosin reporter (Sqh:GFP). Yellow dashes outline the wounds. Time is with respect to wounding. Anterior left, ventral down. Bars, 10 µm. **(C-D)** Wound closure rate constant (C) and mean myosin fluorescence at the wound edge (D) in control (blue, $n$ = 6 wounds) and LIMK-overexpressing embryos (orange, $n$ = 11). **(E)** Kymographs displaying Sqh:GFP FRAP in a segment of the cable at the wound edge in control (top) or LIMK-overexpressing (bottom) embryos. Red asterisks indicate the position of the wound. Yellow boxes show the photobleached region, yellow lines indicate the time of photobleaching. Anterior left, ventral down. Bars, 3 s. **(F-I)** Initial fluorescence before photobleaching (F), percent of fluorescence recovered over time (G), mobile fraction (H), and half-time of fluorescence recovery (I) for FRAP experiments in control ($n$ = 8 wound edge segments) and LIMK-overexpressing embryos ($n$ = 7). (C, F, H-I) Error bars, SD; line, mean. * $P < 0.05$ (Mann-Whitney test). (D, G) Error bars, SEM.

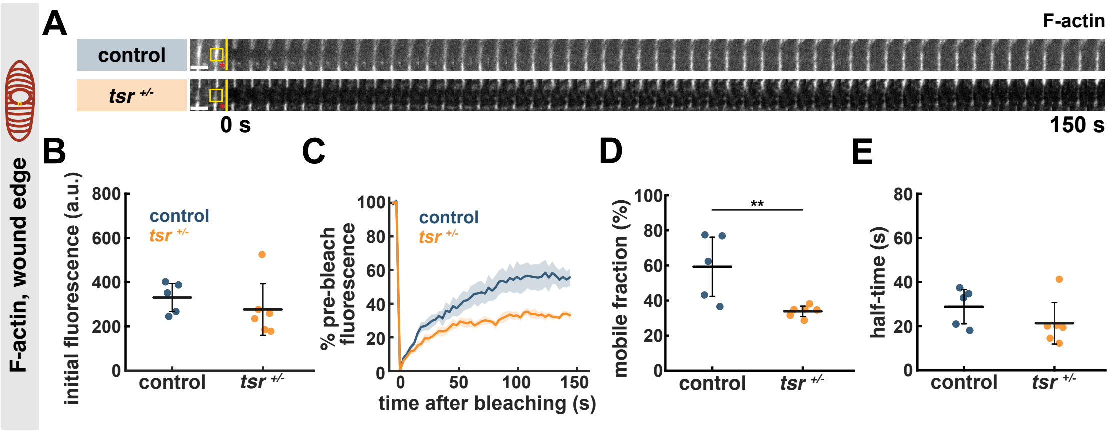


**Figure S4. Cofilin promotes F-actin turnover at the wound edge. (A)** Kymographs displaying GFP:UtrophinABD FRAP in a segment of the cable at the wound edge in control (top) or *tsr* $^{+/-}$ mutant (bottom) embryos. Red asterisks indicate the position of the wound. Yellow boxes show the photobleached region, yellow lines indicate the time of photobleaching. Anterior left, ventral down. Bars, 3 s. **(B-E)** Initial fluorescence before photobleaching (B), percent of fluorescence recovered over time (C), mobile fraction (D), and half-time of fluorescence recovery (E) for FRAP experiments in control (blue, $n = 5$ wounds) or *tsr* $^{+/-}$ mutant embryos (orange, $n = 6$). (B, D-E) Error bars, SD; line, mean. ** $P < 0.001$ (Mann-Whitney test). (C) Error bars, SEM.

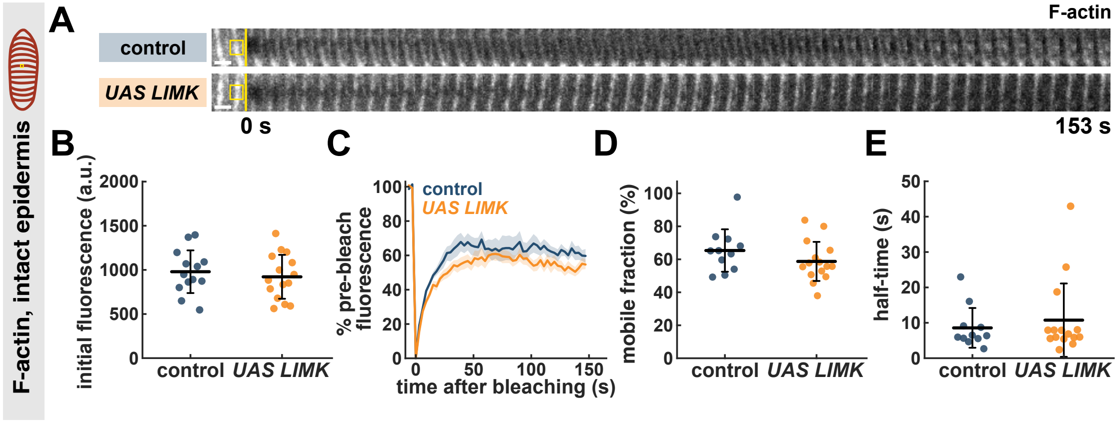


**Figure S5. Cofilin does not control F-actin turnover in the intact epidermis. (A)** Kymographs displaying GFP:UtrophinABD FRAP in a cell-cell junction in the intact epidermis in control (top) or LIMK-overexpressing (bottom) embryos. Yellow boxes show the photobleached region, yellow lines indicate the time of photobleaching. Anterior left, ventral down. Bars, 3 s. **(B-E)** Initial fluorescence before photobleaching (B), percent of fluorescence recovered over time (C), mobile fraction (D), and half-time of fluorescence recovery (E) for FRAP experiments in control (blue, *n* = 13 junctions) or LIMK-overexpressing embryos (orange, *n* = 15). (B, D-E) Error bars, SD; line, mean. (C) Error bars, SEM.

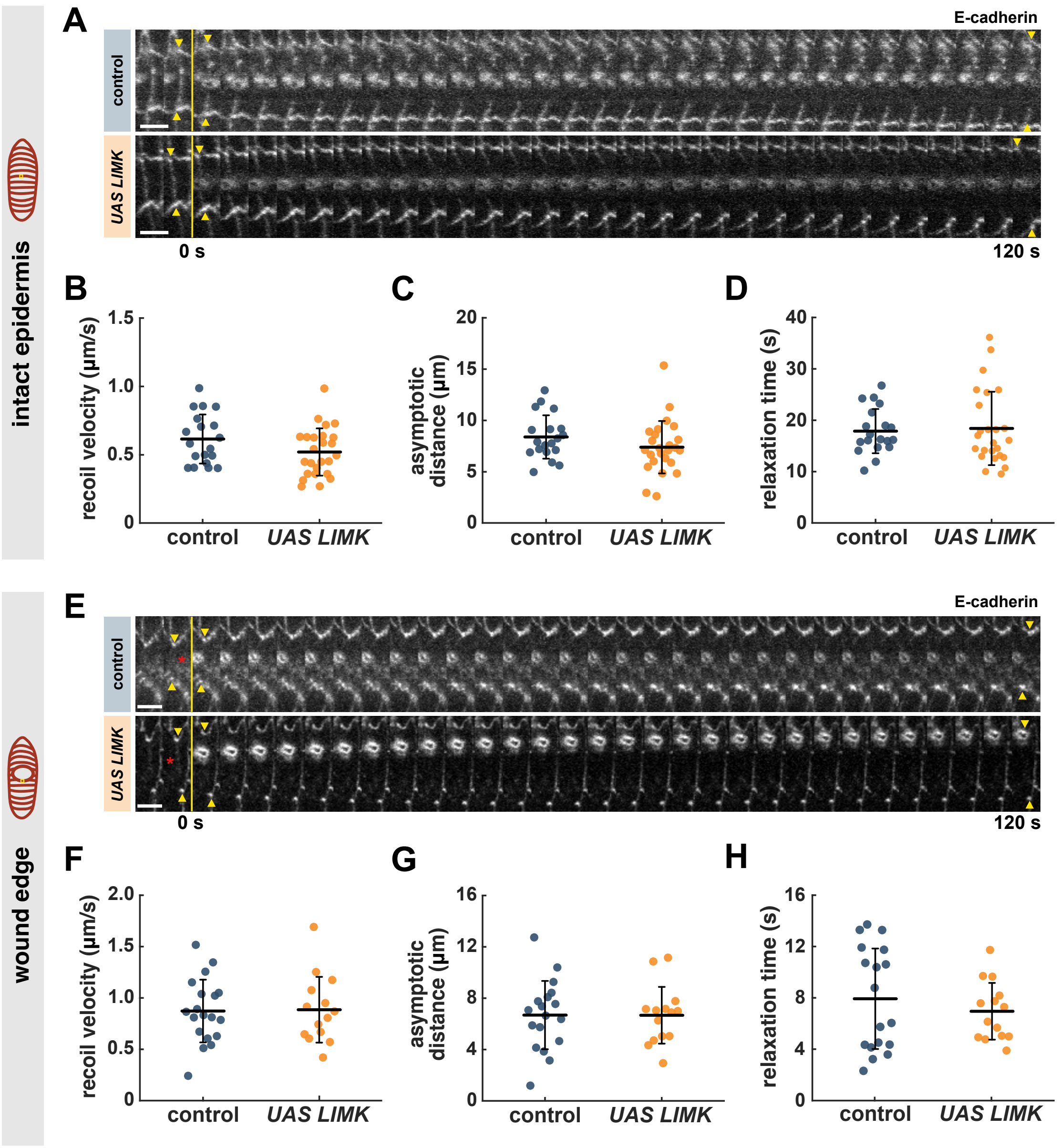


**Figure S6. Cofilin does not control tension or viscoelasticity in the *Drosophila* epidermis. (A, E)** Kymographs showing laser ablation of a cell junction in the intact epidermis (A) or a segment of the wound edge (E) in control (top) or LIMK-overexpressing (bottom) embryos. Arrowheads indicate the ends of the severed structure. Red asterisks indicate the position of the wound. Yellow lines show the time of ablation. Anterior left, ventral down. Bars, 4s. **(B-D, F-H)** Retraction velocity after ablation (B, F), and Kelvin-Voigt model parameters: asymptotic distance retracted (proportional to the tension-to-elasticity ratio, C, G), and relaxation time (proportional to the viscosity-to-elasticity ratio, D, H) in control (blue, $n$ = 19 junctions in B-D, $n$ = 19 wounds in F-H) and LIMK-overexpressing embryos (orange, $n$ = 14 in B-D, $n$ = 25 in F-H). Error bars, SD; line, mean.

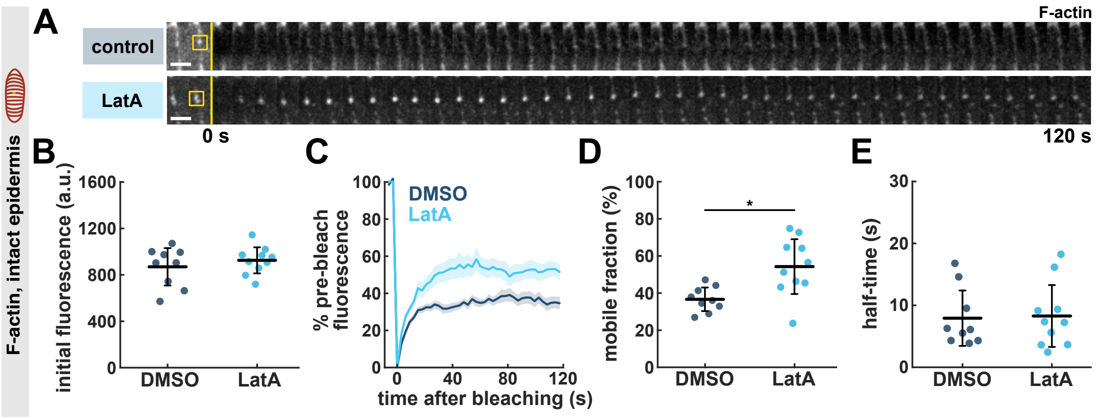


**Figure S7. LatA treatment increases F-actin turnover**. **(A)** Kymographs displaying GFP:UtrophinABD FRAP in a cell-cell junction in the epidermis of embryos treated with 50% DMSO (blue) or 750 µM LatA (cyan). Yellow boxes show the photobleached region, yellow lines indicate the time of photobleaching. Anterior left, ventral down. Bars, 3 s. **(B-E)** Initial fluorescence before photobleaching (B), percent of fluorescence recovered over time (C), mobile fraction (D), and half-time of fluorescence recovery (E) for FRAP experiments in DMSO-treated (blue, $n = 9$ junctions) or LatA-treated (cyan, $n = 10$ junctions) embryos. (B, D-E) Error bars, SD; line, mean. * $P < 0.05$ (Mann-Whitney test). (C) Error bars, SEM.

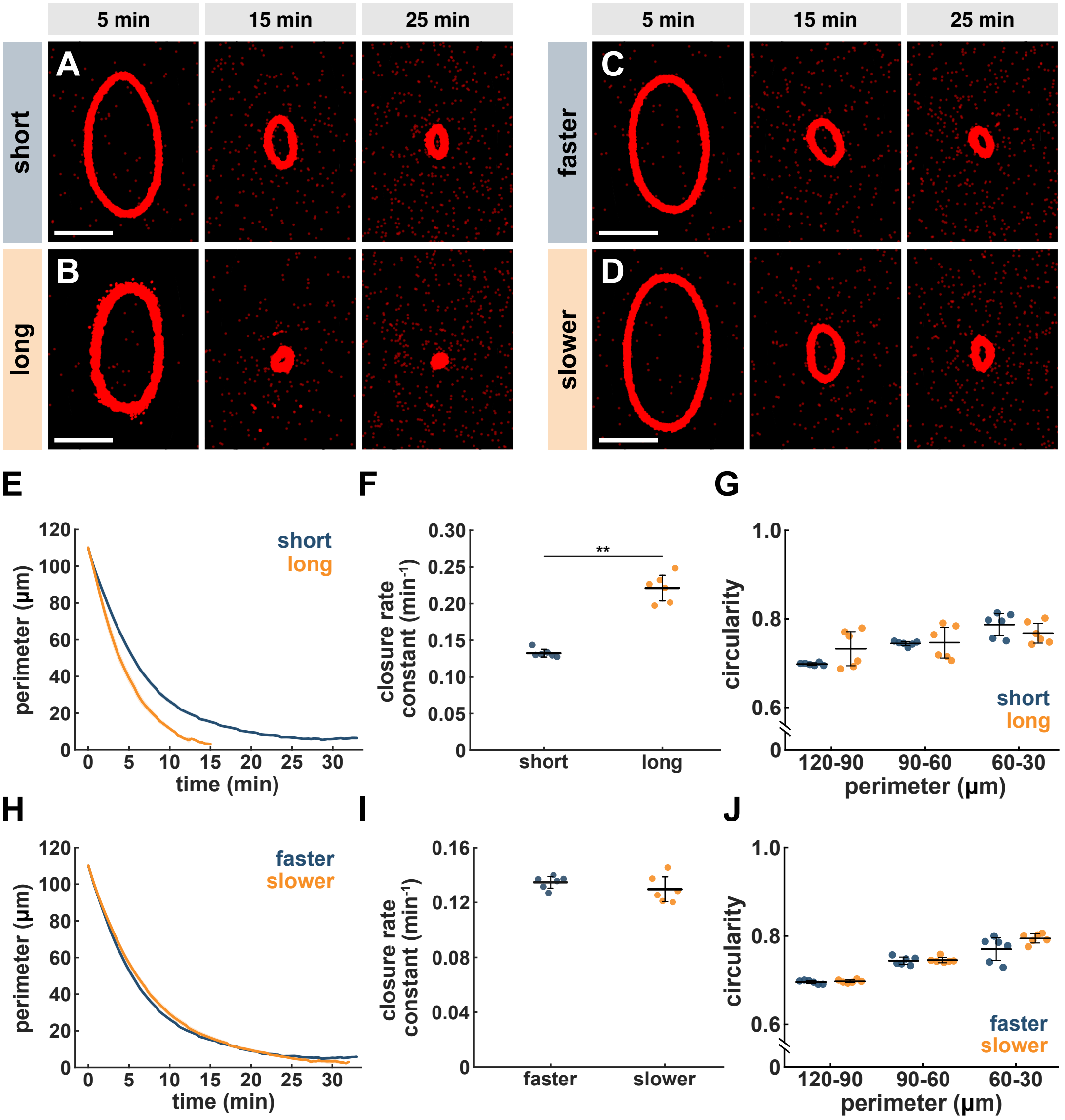


**Figure S8. Computational modelling predicts that changes in filament length and filament disassembly rates do not recapitulate the effects of Cofilin on wound closure. (A-D)** Wound closure in simulations with short (A) or long (B) actin filaments, or with faster (C) or slower (D) F-actin disassembly. Red represents an actin-binding protein to visualize network contraction. Time is with respect to the onset of wound healing. Bars, 10 μm. **(E-J)** Wound perimeter over time (E, H), wound closure rate constant (F, I), and simulation averages of wound circularity binned by wound perimeter (G, J) for simulations with short (blue, $n$ = 6 simulations) or long (orange, $n$ = 6) actin filaments (E-G), and with faster (blue, $n$ = 6) or slower (orange, $n$ = 6) F-actin disassembly (H-J). (E, H) Error bars, SEM. (F-G, I-J) Error bars, SD; line, mean. ** $P < 0.01$ (Mann-Whitney).

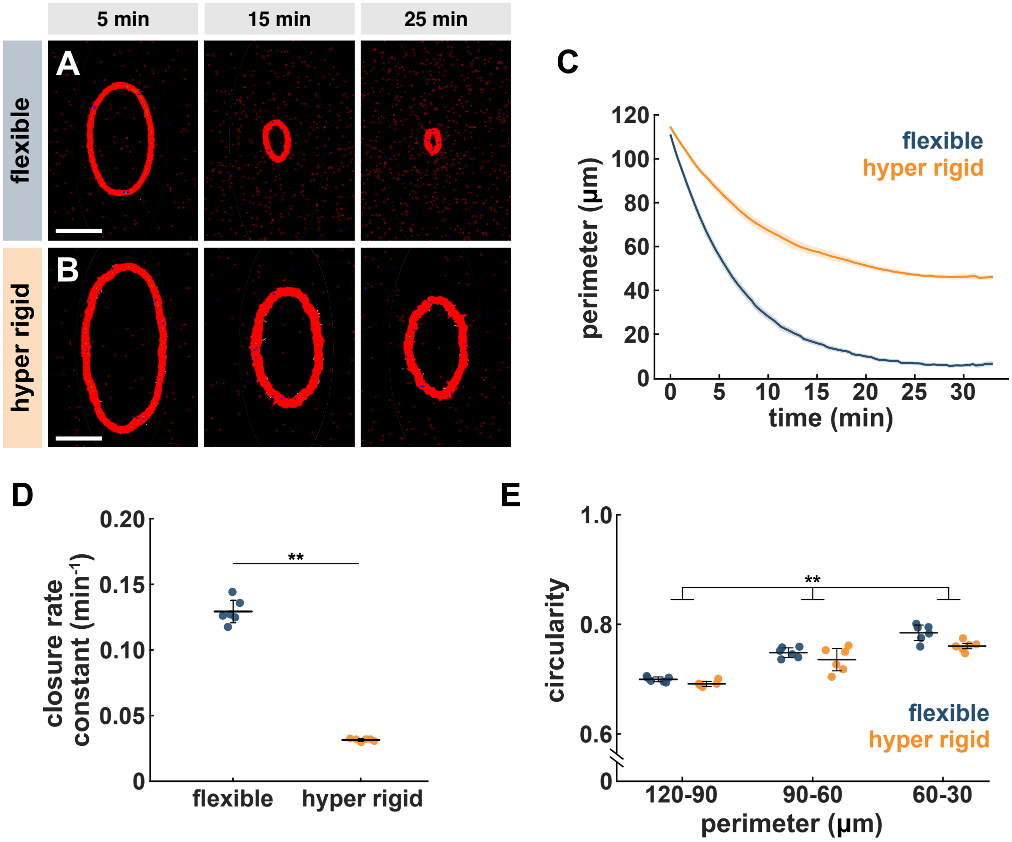


**Figure S9. Computational modelling predicts that the defects in wound closure worsen as F-actin rigidity increases. (A-D)** Wound closure in simulations with flexible (A) or hyper rigid (B) actin filaments. Hyper rigid filaments are 10x more rigid than flexible ones. Red represents an actin-binding protein to visualize network contraction. Time is with respect to the onset of wound healing. Bars, 10 µm. **(C-E)** Wound perimeter over time (C), wound closure rate constant (D), and simulation averages of wound circularity binned by wound perimeter (E) for simulations with flexible (blue, $n$ = 6 simulations) or hyper rigid (orange, $n$ = 6) actin filaments. (C) Error bars, SEM. (D-E) Error bars, SD; line, mean. ** $P < 0.01$ (Mann-Whitney (D), or Fisher's method combining $P$-values for all 3 bins (E)).

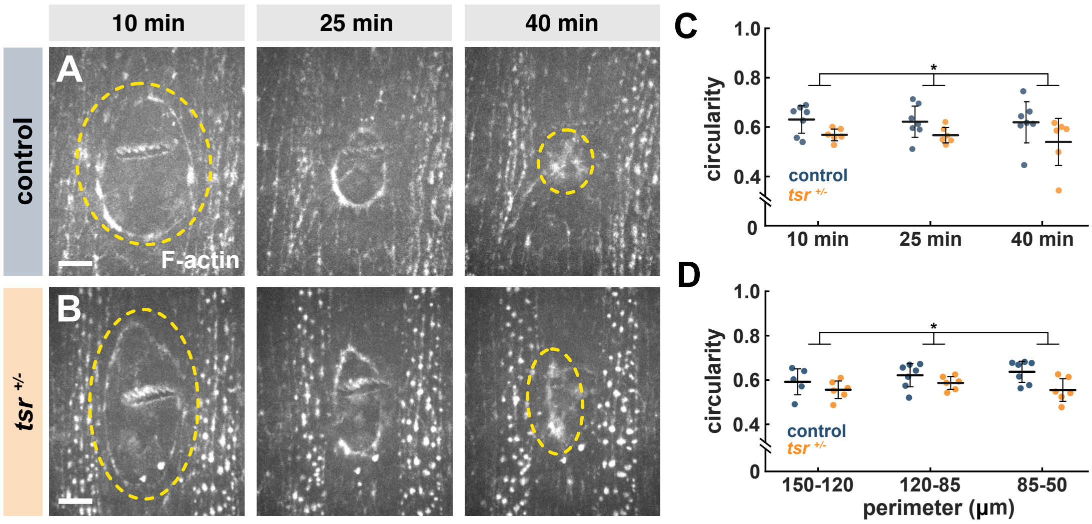


**Figure S10. Cofilin facilitates wound rounding. (A-B)** Wounds at specific times during tissue repair in control (A) and *tsr* $^{+/-}$ mutant (B) embryos. Embryos express GFP:UtrophinABD. Yellow dashes outline the wounds. Anterior left, ventral down. Bars, 10 µm. **(C-D)** Wound circularity at specific times during tissue repair (C), or for wounds of certain perimeter (D) in control (blue, $n$ = 7 wounds) or *tsr* $^{+/-}$ mutant embryos (orange, $n$ = 6). Error bars, SD; line, mean. * $P < 0.05$ (Fisher's method combining *P*-values for all 3 bins).

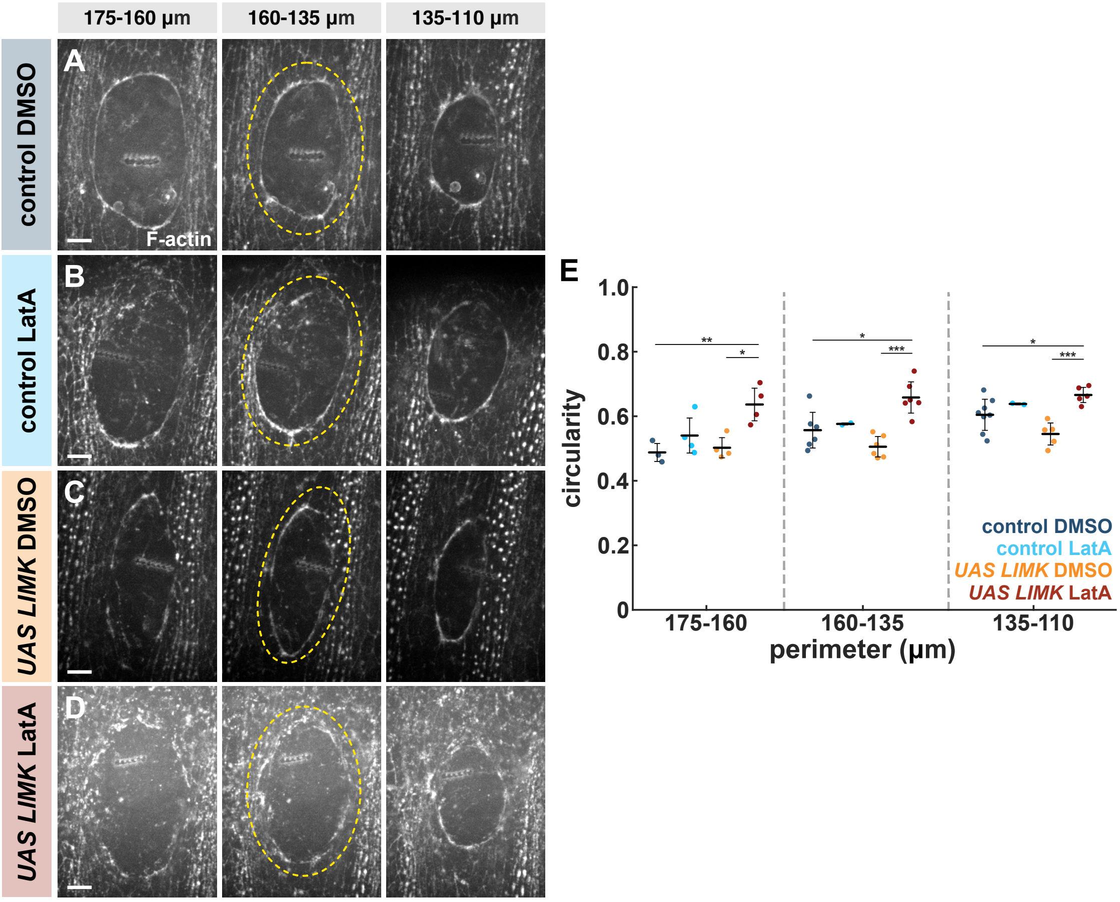


**Figure S11. F-actin turnover facilitates wound rounding**. **(A-D)** Wound closure at different wound perimeters in control (A-B) and LIMK-overexpressing (C-D) embryos treated with 50% DMSO (A, C) or 750 µM LatA (B, D). Embryos expressed GFP:UtrophinABD. Yellow dashes outline the wounds. Anterior left, ventral down. Bars, 10 µm. **(E)** Circularity for wounds of a certain perimeter in control embryos treated with DMSO (blue, $n = 8$ wounds) or LatA (cyan, $n = 3$), and in LIMK-overexpressing embryos treated with DMSO (orange, $n = 6$) or with LatA (red, $n = 5$). Error bars, SD; line, mean. * $P < 0.05$, ** $P < 0.01$, *** $P < 0.001$ (Dunn's test).

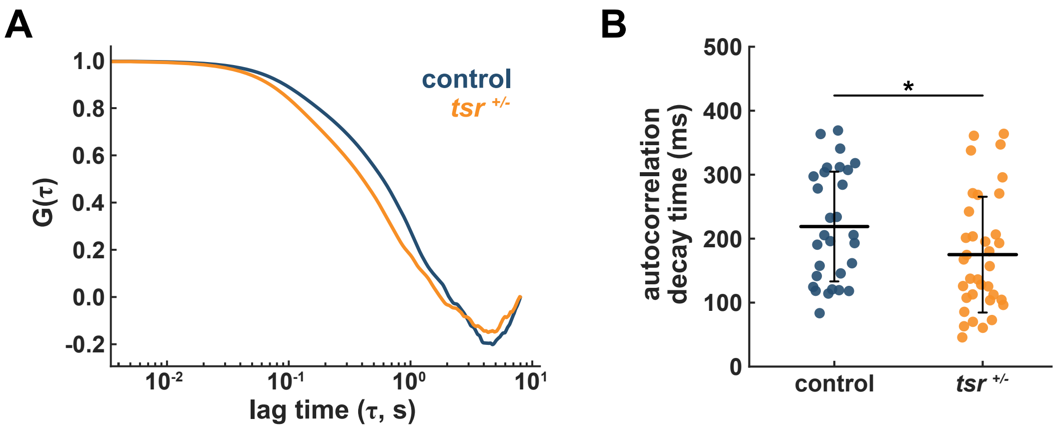


**Figure S12. Cofilin controls F-actin rigidity *in vivo*. (A-B)** Mean autocorrelation, *G* (A) and autocorrelation decay times (B) for GFP:UtrophinABD fluctuations at the wound edge in control (blue, $n = 29$ pinholes in 29 wounds) and *tsr* $^{+/-}$ embryos (orange, $n = 35$ pinholes in 30 wounds). (A) Error bars, SEM. (B) Error bars, SD; line, mean. * $P < 0.05$ (Mann-Whitney test).

**Table S1. Model parameters.** Parameter values used in the simulations when testing the effects on network contraction of severing, filament length, filament rigidity, and filament disassembly rate.

| parameter | severing | length | rigidity | disassembly | source |
|---|---|---|---|---|---|
| time step | 0.1 s | 0.1 s | 0.1 s | 0.1 s | |
| crosslinker length | 0.1 μm | 0.1 μm | 0.1 μm | 0.1 μm | (default) |
| crosslinker stiffness | 100 pN/μm | 100 pN/μm | 100 pN/μm | 100 pN/μm | (default) |
| actin filaments per bundle | 8 | 8 | 8 | 8 | (calibration) |
| crosslinkers per bundle | 40 | 80 | 40 | 40 | (calibration) |
| bundles per network | 1000 | 500 | 1000 | 1000 | (calibration) |
| # of myosin minifilaments | 1600 | 1600 | 1600 | 1600 | (calibration) |
| myosin minifilament rigidity | 1 pN*μm$^2$ | 1 pN*μm$^2$ | 1 pN*μm$^2$ | 1 pN*μm$^2$ | (default) |
| motor binding rate | 5 $s^{-1}$ | 5 $s^{-1}$ | 5 $s^{-1}$ | 5 $s^{-1}$ | (default) |
| motor dissociation rate | 1 $s^{-1}$ | 1 $s^{-1}$ | 1 $s^{-1}$ | 1 $s^{-1}$ | (default) |
| # of severing molecules | 0 | 20 | 20 | 20 | (calibration) |
| actin filament rigidity | 0.01 pN*μm$^2$ | 0.01 pN*μm$^2$ | 0.04 pN*μm$^2$ | 0.01 pN*μm$^2$ | (McCullough et al., 2008) |
| actin filament length | 3 μm | 6 μm | 3 μm | 3 μm | (Robaszkiewicz et al., 2020) |
| actin disassembly rate | 0.001 μm/s | 0.001 μm/s | 0.001 μm/s | 0.0005 μm/s | (calibration) |
| actin assembly rate | 0 μm/s | 0 μm/s | 0 μm/s | 0 μm/s | (default) |
| initial circularity | 0.75 | 0.75 | 0.75 | 0.75 | (measured) |

### Supplementary video legends

**Video S1. Cofilin is necessary for rapid wound closure.** Wound closure in control (left) and LIMK-overexpressing (right) embryos expressing GFP:UtrophinABD. Images were acquired every 30 s. Time after wounding is shown. Anterior left, ventral down.

**Video S2. Precise levels of F-actin turnover are necessary for rapid wound healing.** Wound closure in control (first and second from the left) or LIMK-overexpressing embryos (third and fourth) injected with 50% DMSO (first and third) or 750 µm of LatA (second and fourth). Images were acquired every 30 s. Time after wounding is shown. Anterior left, ventral down.

**Video S3. Computational modelling predicts that Cofilin may induce F-actin severing to drive rapid wound repair.** Wound closure in simulations with (left) or without (right) F-actin severing. Red represents an actin-binding protein to visualize network contraction. Images were exported every 20 s. Time is with respect to the onset of wound healing.

**Video S4. Computational modelling predicts that Cofilin may control F-actin rigidity to drive rapid wound repair.** Wound closure in simulations with flexible (left) or rigid (right) F-actin. Red represents an actin-binding protein to visualize network contraction. Images were exported every 20 s. Time is with respect to the onset of wound healing.
